# Loss of the Alzheimer’s-linked bridging integrator 1 (BIN1) protein affects synaptic structure and disrupts tau localisation and release

**DOI:** 10.1101/646406

**Authors:** Elizabeth B. Glennon, Dawn H-W Lau, Rebecca M.C. Gabriele, Matthew F. Taylor, Claire Troakes, Christina Elliott, Richard Killick, Diane P. Hanger, Beatriz G. Perez-Nievas, Wendy Noble

**Author notes:** **Corresponding Authors:** Dr Elizabeth Glennon, King’s College London, Institute of Psychiatry, Psychology and Neuroscience, Department of Basic and Clinical Neuroscience, Maurice Wohl Clinical Neuroscience Institute, 5 Cutcombe Road, London, SE5 9RX. UK. Dr Wendy Noble, King’s College London, Institute of Psychiatry, Psychology and Neuroscience, Department of Basic and Clinical Neuroscience, Maurice Wohl Clinical Neuroscience Institute, Rm1.23, 5 Cutcombe Road, London, SE5 9RX. UK.

## Abstract

**Background:** Post-translational modifications of tau modify its interaction with binding partners and cause tau mislocalisation and altered tau function in Alzheimer’s disease (AD). The AD risk gene BIN1, is a binding partner for tau, however the mechanism by which BIN1 influences tau function is not fully understood. We hypothesised that BIN1 modulates AD risk by causing damaging tau mis-sorting to the synapse.

**Methods:** Tau and BIN1 levels, distribution and interactions were assessed in post-mortem control and AD brain and in primary neurons. In primary neurons, tau was further examined using structured illumination microscopy and immunoblotting following BIN1 knockdown, BIN1-tau interactions were examined using proximity ligation assays and tau release from neurons was measured by sensitive sandwich ELISA.

**Results:** Proline 216 in tau was identified as critical for tau interaction with the BIN1-SH3 domain, and tau phosphorylation at serine/threonine residues disrupted this interaction. Subcellular fractionation showed that BIN1 is lost from the cytoplasm of AD brain and this correlated with the mislocalisation of phosphorylated tau to synapses. Mimicking BIN1 loss in AD by knockdown of the protein in primary neurons altered the structure of dendritic spines, caused phosphorylated tau to mis-sort to synapses and reduced the physiological release of predominantly dephosphorylated tau.

**Conclusions:** These data suggest that BIN1 loss in AD allows phosphorylated tau to be mis-sorted to synapses which likely alters the integrity of the post-synapse, alongside reducing the functionally important release of physiological forms of tau.

## Introduction

Genome-wide association studies (GWAS) identified variants of bridging integrator 1 (*BIN1*) that confer the second largest genetic risk factor for developing sporadic AD, after APOE4 (1–7). BIN1 is a cytoplasmic membrane-binding protein, which plays important roles in endocytosis and subcellular trafficking (8). BIN1 has a number of splice variants with tissue specific expression (8). The more common *BIN1* variants are upstream of the gene and do not affect protein structure but may affect splicing or expression of BIN1. In AD brain, expression of the longer neuronal forms of BIN1 are decreased, while the shorter glial isoforms are increased (9–11). Chapuis *et al.* (12) recently reported a 3 base pair insertion in complete linkage disequilibrium with the BIN1 polymorphism rs744373 that is associated with increased mRNA levels of BIN1 (12), although Adams *et al.* (13) found no correlation between rs744373 and BIN1 expression. Nevertheless, BIN1 risk polymorphisms at rs744373 are associated with increased BIN1 expression and poorer memory in those with epilepsy (14, 15).

Alterations to tau protein are closely associated with synaptic dysfunction, neurodegeneration and cognitive decline in AD (16, 17). Direct interactions between BIN1 and tau have been demonstrated in cell models, mice and Drosophila (12, 18–20), BIN1 colocalises with neurofibrillary tangles (11) and is associated with elevated tau phosphorylation in AD brain (21). Importantly, expression of BIN1 in a Drosophila model of AD was recently shown to modulate the toxicity of tau (12). Others have shown that BIN1 over-expression in mice causes microstructural changes in hippocampal circuits (22). These are the first circuits to show tau pathology in AD (22), suggesting that BIN1 may affect the development of AD by modulating tau effects at synapses and possibly also synaptic activity-dependent tau release (23). Therefore, the purpose of this study was to investigate the functional consequences of interactions between BIN1 and tau for disease-associated tau mis-sorting and release and synapse health.

## Materials and methods

### Human Brain

Braak staged post-mortem human brain tissue was obtained from the MRC London Neurodegenerative Diseases brain bank at King’s College London following ethical approval (REC reference: 08/MRE09/38 + 5). Details of these samples are provided in Tables 1 and 2.

**Table 1:**
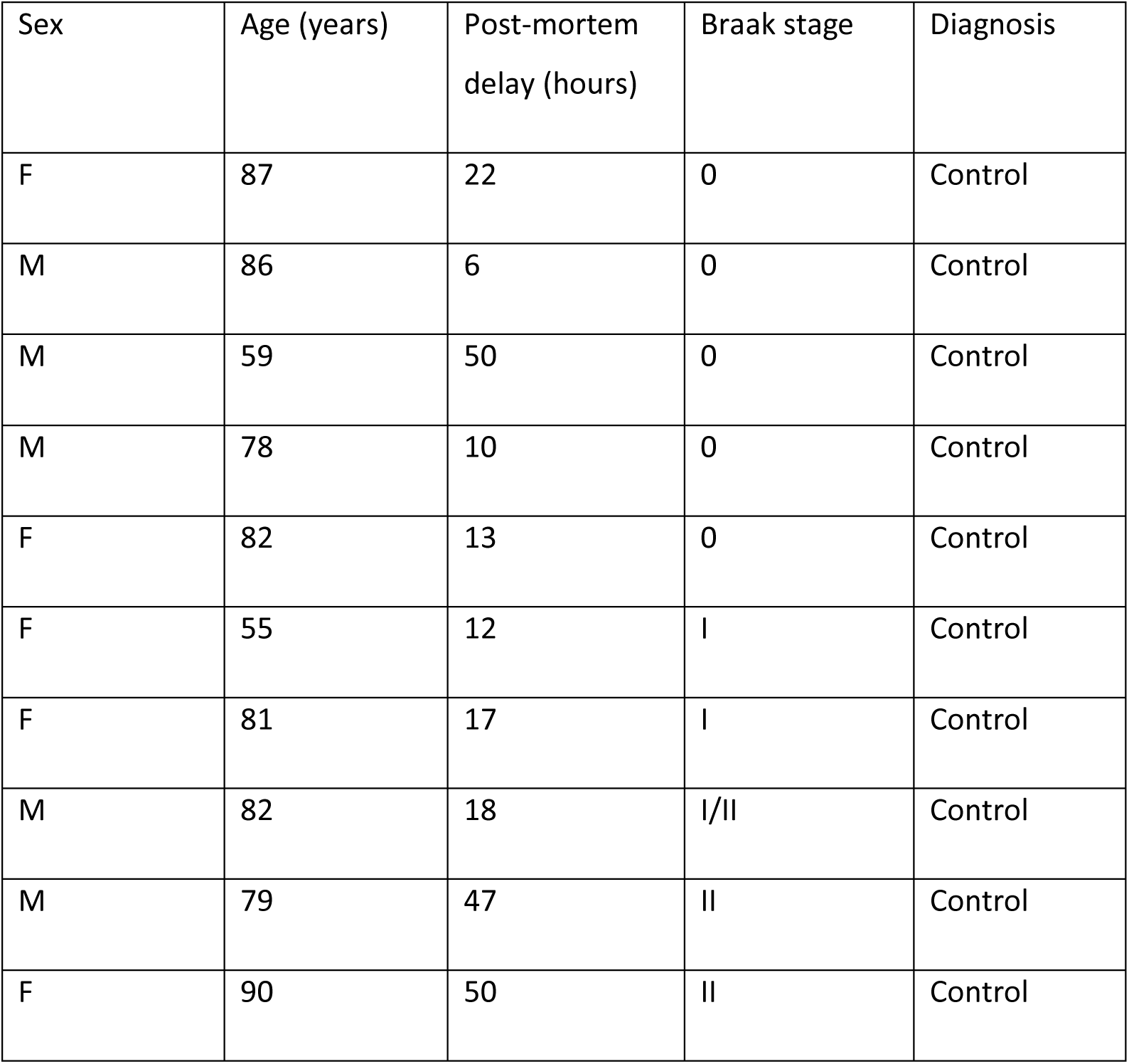
Characteristics of human brain tissue (paraffin embedded frontal cortex sections) used for proximity ligation assays. Table shows details of sex, age, post-mortem delay (hours), Braak stage and AD diagnosis for cases from which paraffin embedded frontal cortex sections were used in this study.

| Sex | Age (years) | Post-mortem delay (hours) | Braak stage | Diagnosis |
| --- | --- | --- | --- | --- |
| F | 87 | 22 | 0 | Control |
| M | 86 | 6 | 0 | Control |
| M | 59 | 50 | 0 | Control |
| M | 78 | 10 | 0 | Control |
| F | 82 | 13 | 0 | Control |
| F | 55 | 12 | I | Control |
| F | 81 | 17 | I | Control |
| M | 82 | 18 | I/II | Control |
| M | 79 | 47 | II | Control |
| F | 90 | 50 | II | Control |

**Table 2:** Characteristics of temporal cortex tissue used in this study. Table shows details of sex, age, post-mortem delay (hours), Braak stage and AD diagnosis for cases from which frozen temporal cortex sections was obtained.

| Sex | Age (years) | Post-mortem delay (hours) | Braak stage | Diagnosis |
| --- | --- | --- | --- | --- |
| F | 74 | 64 | II | Control |
| F | 90 | 44 | II | Control |
| F | 73 | 27 | I | Control |
| F | 77 | 21 | 0 | Control |
| F | 80 | 22 | II | Control |
| M | 68 | 60 | II | Control |
| M | 80 | 55 | II-III | Control |
| M | 90 | 45 | - | Control |
| M | 78 | 24 | III | Control |
| F | 92 | 9 | II | Control |
| M | 82 | 47 | I | Control |
| F | 84 | 34 | I-II | Control |
| F | 90 | 50 | II | Control |
| M | 66 | 52 | - | Control |
| M | 82 | 18 | I/II | Control |
| M | 91 | 48 | IV | Moderate AD |
| M | 88 | 79 | III-IV | Moderate AD |
| F | 95 | 47 | IV | Moderate AD |
| M | 84 | 86 | IV | Moderate AD |
| M | 98 | 53 | IV | Moderate AD |
| F | 86 | 55.5 | IV | Moderate AD |
| M | 82 | 28 | IV | Moderate AD |
| M | 86 | 52.5 | IV | Moderate AD |
| F | 83 | 22 | IV | Moderate AD |
| M | 93 | 13.5 | IV | Moderate AD |
| F | 83 | 41.5 | IV | Moderate AD |
| F | 97 | 67.5 | III-IV | Moderate AD |
| F | 96 | 39 | IV | Moderate AD |
| F | 92 | 19.5 | III | Moderate AD |
| F | 92 | 29.5 | IV | Moderate AD |
| F | 73 | 30 | VI | Severe AD |
| F | 84 | 27 | VI | Severe AD |
| F | 79 | 31 | VI | Severe AD |
| M | 86 | 38 | VI | Severe AD |
| F | 85 | 79 | VI | Severe AD |
| M | 67 | 39.5 | VI | Severe AD |
| F | 69 | 73 | VI | Severe AD |
| F | 89 | 38.5 | VI | Severe AD |
| F | 93 | 49 | VI | Severe AD |
| M | 84 | 67 | VI | Severe AD |
| F | 81 | 20 | VI | Severe AD |
| M | 83 | 22 | VI | Severe AD |
| F | 81 | 17.5 | VI | Severe AD |
| F | 86 | 25 | VI | Severe AD |
| M | 66 | 41 | VI | Severe AD |

### Modification of BIN1 expression in primary neurons

Primary cortical neurons were dissected from E18 Sprague-Dawley rats and cultured as previously described (24) on poly-D-lysine coated plates or glass cover-slips. Lentivirus shRNA targeting BIN1 from the RNAi consortium (TRCN0000088188) and a scrambled control sequence in the pLKO.1 vector were purchased from Dharmacon Horizon (CO, USA). PAX2 and pMG.2 plasmids were kind gifts from Dr Maria Jimenez-Sanchez (King’s College London). HEK293 cells cultured in Dulbecco’s Modified Eagle medium plus GlutaMAX (DMEM, Thermo Fisher Scientific, MA, USA) supplemented with 10 % (v/v) fetal bovine serum (FBS, Thermo Fisher Scientific) were transfected with PAX2, pMG.2, and shRNA using Lipofectamine 2000 (Invitrogen, CA, USA). Lentiviral particles collected from culture medium were isolated and concentrated according to the manufacturer’s instructions. BIN1 was knocked down using either lentiviral shRNA constructs or Accell siRNA smart pool (E-095528, Dharmacon Horizon). At 19 days *in vitro* (DIV), rat primary cortical neurons were transfected with 50 nM BIN1 or non-targeting control siRNA (Dharmacon Horizon Discovery) using Lipofectamine 2000 and incubated for 96 hours at 37°C before use. For lentiviral knockdown, neurons were cultured for 5 DIV, then treated with either BIN1 targeting shRNA or scrambled control shRNA lentiviral particles for 24 hours, after which time the virus was removed and neurons further cultured until 21-23 DIV.

### GST binding assays

BIN1-SH3 cDNA generously provided by Isabelle Landrieu (University of Lille Nord de France) was cloned into pGEX5X1 using sequence and ligation independent cloning (SLIC) (25). The BIN1-SH3 domain was amplified from the original vector using primers 5’-TCG AGC GGC CGC ATC GTG ACA TGG GTC GTC TGG ATC TG-3’ and 5’-AAA CGC GCG AGG CAG ATC GTC AGT TAC GGC ACA CGC TCA GTA AAA TTC-3’, and pGEX5X1 was linearized using primers 5’-CTG ACG ATC TGC CTC GCG-3’ and 5’-GTC ACG ATG CGG CCG CTC-3’. SLIC products were used to transform BL21 *E.coli* (New England Biolabs, MA, USA) by heat shock. DNA was purified using QIAgen spin miniprep kit (QIAgen, Hilden, Germany) and the cloning was confirmed by sequencing (Source Bioscience, Nottingham, UK, using stock primers to GST plasmid). BL21 *E.coli* containing either BIN1-SH3-pGEX5X1 or empty vector pGEX5X1 were used to produce glutathione S-transferase (GST) fusion proteins and GST-pulldown assays were performed as described previously (26). WT and PxxP mutant tau plasmids have been described previously (26). These were expressed in HEK293 cells for 24 hours after which time cells were lysed and the lysates used in GST pull-downs.

### Tau ELISA and cell viability assays

Lactate dehydrogenase was measured in culture medium to determine cell health, and sandwich ELISAs were used to determine tau content in culture medium from neurons incubated for 4 hours in Hank’s balanced salt solution (HBSS), without Ca^2+^ and Mg^2+^ as described previously (27).

### Proximity Ligation Assays

23 DIV neurons were fixed in 4 % (w/v) paraformaldehyde in phosphate-buffered saline (PBS) (Alfa Aesar, MA, USA) and non-specific fluorescence quenched by incubation in 50 mM NH_4_Cl in PBS for 10 minutes, followed by cell permeabilisation in 0.2 % (v/v) Triton X-100 in PBS. Proximity ligation assays (PLAs) were performed using Duo-link reagents (Sigma Aldrich, MO, USA), according to the manufacturer’s protocol and with primary antibodies against BIN1 (ab54764, Abcam, Cambridge, UK) and total tau (Agilent, CA, USA). Actin was labelled with phalloidin-488 (Life Technologies, CA, USA), and nuclei stained with 10 μgml^−1^ Hoescht-33352 (Thermo Fischer Scientific). Image stacks covering the whole volume of each cell were acquired using a Nikon Eclipse Ti-E microscope and analysed using Fiji. Z-stacks were converted to maximum intensity projections, and the phalloidin signal used to identify cell outlines. The intensity of PLA signals in each cell was quantified using Fiji (28) and is expressed as a proportion of cell area.

PLAs on paraffin-embedded human frontal cortex were performed as described above, with the following additions. Sections were rehydrated and de-waxed using standard methods, and antigen retrieval was performed by 1-hour incubation in 10 mM glycine, 0.25 % (v/v) Triton X-100 in Tris-buffered saline (TBS). Sections were blocked for two hours in 5 % (v/v) horse serum, 0.25 % (v/v) Triton X-100 in TBS. MAP2 (GTX82661, GeneTex, CA, USA) was immunolabelled and detected using Alexa Fluor-conjugated secondary antibodies (Life Technologies).

### Immunofluorescence

Immunofluorescence was performed as described (29), but using 2 % (v/v) fetal bovine serum (FBS, Life Technologies). Cells were incubated with primary antibodies against BIN1 (ab54764, Abcam), PSD-95 (D74D3, Cell Signalling), synaptophysin (sc7568, Santa Cruz, TX, USA) and MAP2 (GTX82661, GeneTex), and the appropriate species of AlexaFluor-conjugated secondary antibody (Life Technologies). Labelled proteins were imaged using an Eclipse Ti2 inverted Nikon 3D N-SIM microscope and images reconstructed using NIS elements software.

### Analysis of synapses

Neurons were fractionated to generate a synapse-enriched fraction using a protocol modified from (30). Synaptoneurosomes were isolated from post-mortem temporal cortex as described in (31, 32). Equal protein concentrations of total, synaptic and cytoplasmic fractions were immunoblotted for tau, BIN1, synaptic and cytoplasmic markers.

Neurons at 22 DIV were transfected with an eGFP-N2 plasmid (Clontech, Kyoto, Japan) using Lipofectamine 2000 for 24 hours, fixed as described above and imaged using a Nikon Eclipse Ti-2 inverted microscope with Vt-iSIM scan head, 3×3 large image stacks were acquired covering the entire volume of the neuron, with 0.2 μm between each image in the Z plane. Neurolucida™ software (MBF Bioscience, VT, USA) was used to trace neurons and detect, classify and quantify dendritic spines.

### SDS-PAGE and western blotting

Samples were electrophoresed on 10 % Tris-glycine-Sodium Dodecyl Sulphate-polyacrylamide gels, Nu-Page 4-12 % or 10 % Bis-Tris gels (Invitrogen), transferred to 0.45 μm nitrocellulose membrane (Millipore, MA, USA) and immunoblotted as described (10). Primary antibodies were BIN1 (99D, Millipore), GST (GE Healthcare, IL, USA), total tau (total human tau, Agilent), Tau-1 (Millipore), PHF1 (Peter Davies, Donald and Barbara Zucker School of Medicine at Hofstra, Northwell, USA), β-actin (ac15, abcam), synaptophysin (sc17750, Santa Cruz), and post-synaptic density-95 (PSD95) (MAB 1596, Millipore). Horseradish peroxidise (HRP)-conjugated secondary antibodies (GE Healthcare) were detected using enhanced chemiluminescence (Thermo Fisher Scientific) and visualised using a Chemi-Doc imager (Bio-Rad, CA, USA). Densitometric analysis was performed using Fiji (28).

### Peptide Arrays

Tau peptide arrays were generated as described previously (33). To determine the BIN1-SH3 domain binding sites on human 2N4R tau, 50 μg recombinant BIN1 in RIPA buffer (50 mM Tris-HCl, 150 mM NaCl, 0.5 % (w/v) sodium deoxycholate, 1 % (v/v) NP-40, pH 8.0) supplemented with ethylenediaminetetraacetic acid and cOmplete protease inhibitor cocktail (Roche Diagnostics, Risch-Rotkreuz, Switzerland), was gently shaken with the peptide array overnight at 4°C. Following washing, arrays were probed with anti-BIN1 antibody (99D, Millipore) and HRP-conjugated secondary antibodies used to detect binding using enhanced chemiluminescence (Thermo Fisher Scientific).

### Data analysis and statistics

Statistical tests were performed using GraphPad Prism 7.0 (CA, USA) or RStudio.

## Results

### Tau and BIN1 interact

BIN1 and tau co-immunoprecipitate following over-expression of both proteins in cell lines, and the endogenous proteins interact in synaptoneurosomes isolated from mouse brain (12). Using a combination of GST-pull downs and NMR spectroscopy Sottejeau *et al.* (20) demonstrated that the BIN1-SH3 domain interacts with the proline-rich region of tau. To extend these findings, we generated GST-BIN1-SH3 constructs (Supplementary Figure 1A) and used these together with GST-only constructs in binding assays with lysates from HEK293 cells transfected with human 2N4R tau or empty vector. Immunoblotting of pull-downs with an antibody against total (non-phosphorylated and phosphorylated) tau confirmed that the BIN1-SH3 domain binds human 2N4R tau (Figure 1A). No interactions were observed in GST-only and empty vector controls. Binding between endogenous BIN1 and tau was confirmed in 18-23 DIV rat primary cortical neurons by proximity ligation assay (PLA). Specificity of the signal was confirmed by siRNA-mediated BIN1 knockdown, which reduced BIN1 expression by approximately 83 % without affecting tau levels (Figure 1B). Strong PLA signals identified in the soma of neurons (Figure 1C) were markedly reduced upon BIN1 siRNA knockdown (Figure 1D); residual interactions likely occurring as a result of incomplete knockdown. Super-resolution N-SIM microscopy also showed distinct regions of tau-BIN1 interactions in the neuronal processes of these cells. PLA signals were closely apposed to but did not co-localise with phalloidin, suggesting that although both BIN1 and tau are actin binding proteins (34–36), their interactions in primary neurons do not take place on the actin cytoskeleton (Figure 1E). To demonstrate that BIN1 interacts with tau in human brain, PLA was also performed on human frontal cortex sections from non-cognitively impaired controls. Strong PLA signals were detected in tissue sections probed with both BIN1 and tau antibodies, but not in the negative control lacking primary antibodies (Figure 1F).

**Figure 1:**
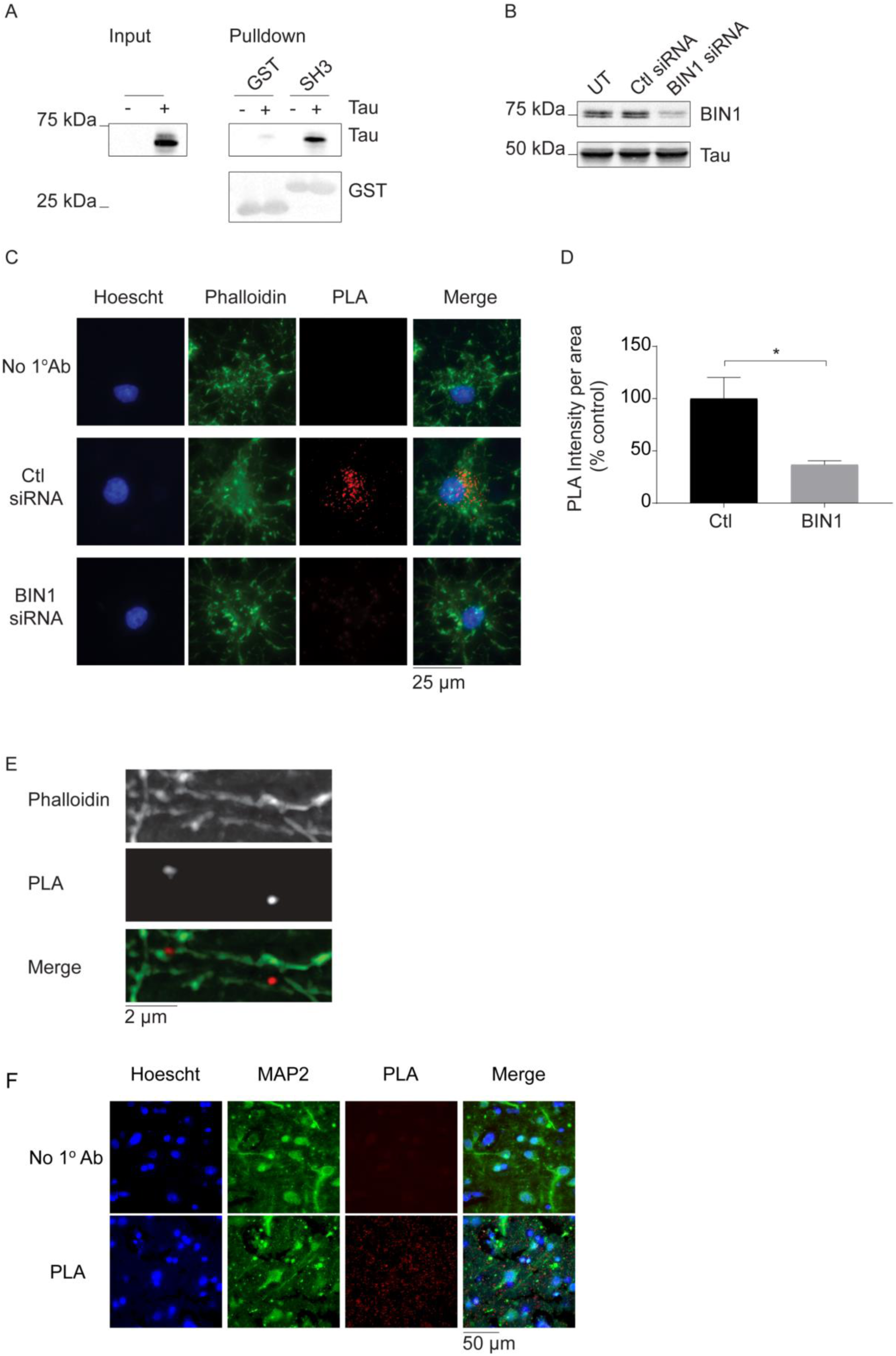
Tau interacts with BIN1. A) Lysates from HEK293 cells transfected with empty vector (−) or 2N4R human tau (+) were incubated with BIN1-SH3-GST glutathione beads, or GST-beads as a control. Lysates (input) and GST-bound proteins (pulldown) were probed on western blots with antibodies against total (phosphorylated and nonphosphorylated) tau (top) or GST (bottom). Tau was pulled down by BIN1-SH3-GST but not GST-only. B) Cell lysates from primary cortical neurons transfected with either non-targeting control siRNA (ctl siRNA) or BIN1 targeting siRNA were western blotted with antibodies against BIN1 (top) and tau (bottom) and show reduced BIN1 protein following treatment with BIN1 siRNA. C) Proximity ligation assays (PLA) were used to demonstrate interactions between endogenous BIN1 and tau in rat primary cortical neurons and to confirm BIN1 knockdown by BIN1 siRNA. Images show PLA signals (red) in neurons treated with Ctl (non-targeting) siRNA indicating regions of BIN1-tau interactions. Nuclei of cells were stained with Hoechst 33352 (blue) and the actin cytoskeleton labelled with phalloidin (green). No PLA signals were observed in controls lacking primary antibody or in primary cortical neurons treated with BIN1 siRNA. D) Bar chart shows significantly reduced PLA signal intensity/area in neurons exposed to BIN1 siRNA (BIN1) relative to non-targeting control (Ctl) siRNA. Data were analysed using Mann-Whitney tests. Data shown are mean ±S.E.M. n=3, *p<0.05. E) N-SIM super resolution imaging of PLA signal (middle) and the actin cytoskeleton (top) in processes of cortical neurons. BIN1 interacts with tau in the vicinity of actin but interactions are not directly associated with the actin cytoskeleton. F) PLA (red) showing interactions between BIN1 and tau in post-mortem human temporal cortex. Nuclei were stained with Hoescht-33352 (blue), and neurons immunolabelled with an antibody against MAP2 (green). No PLA signals were obtained in controls omitting primary antibody.

### BIN1 binds to P216 in the proline-rich region of tau and this interaction is regulated by tau phosphorylation

Tau peptide arrays (33), were incubated with 50 µg recombinant BIN1-SH3-GST, GST, or 2 % BSA, and probed with a BIN1 antibody. This yielded a strong signal in the BIN1-SH3-GST array only for the peptide corresponding to amino acids 201-225 in the proline-rich domain of 2N4R tau (peptide 41 in Figure 2A, B, Supplementary Figure 2A). PxxP sequences, of which there are seven in the proline-rich region of tau (26), are the minimum consensus target sites for SH3 domain binding. To identify which PxxP motifs are required for the BIN1-tau interaction, we performed BIN1-SH3-GST pulldown assays following transfection of HEK293 cells with WT 2N4R tau or constructs in which each tau PxxP motif (Supplementary Figure 2B) was disrupted by mutating a single proline to alanine (26, 37). There was a significant reduction in BIN1-SH3-GST binding to P216A relative to WT tau (Figure 2C, D), demonstrating that proline 216 in tau is critical for the tau-BIN1 interaction.

**Figure 2:**
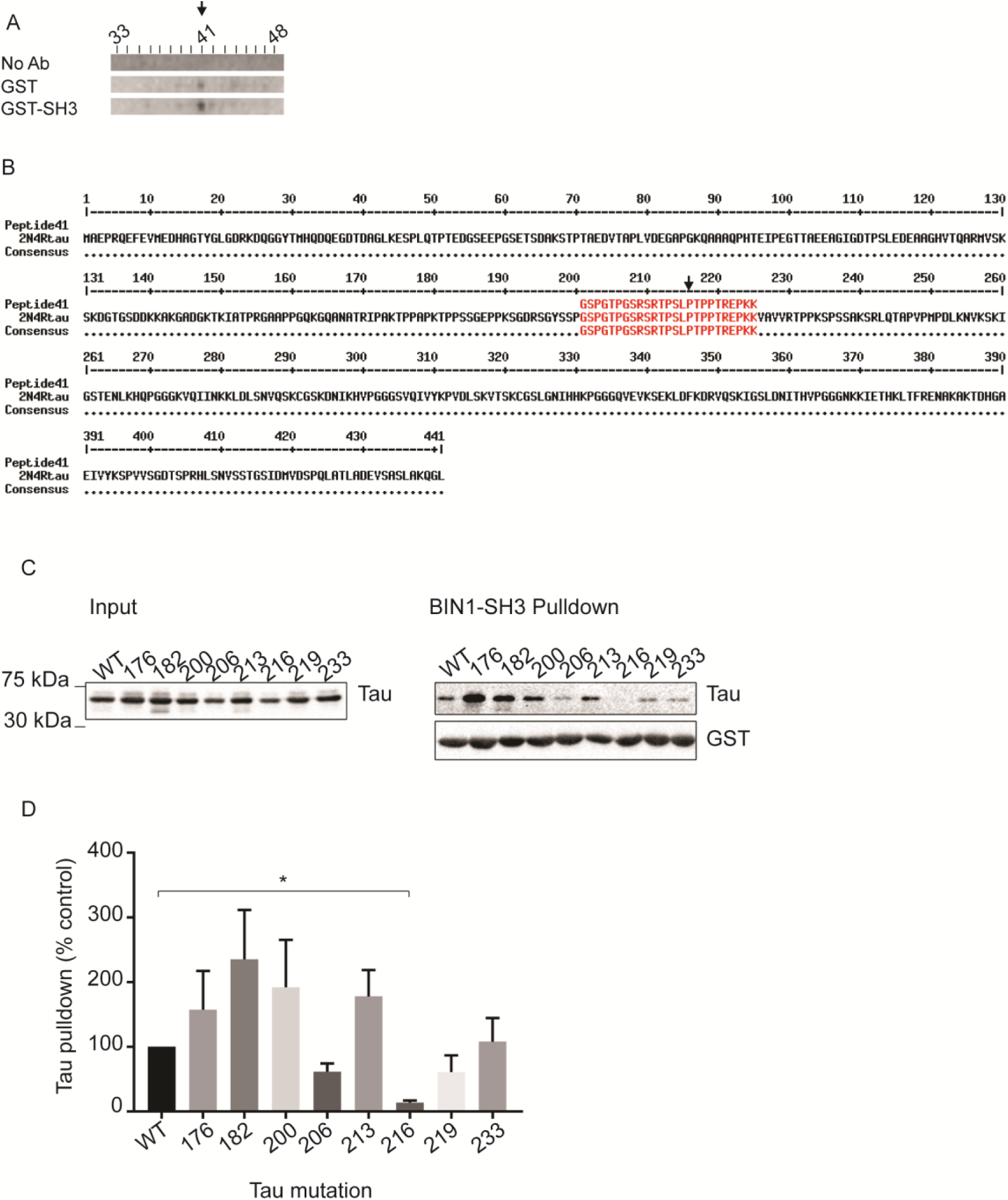
The BIN1-SH3 domain binds to PxxP motifs in tau. A) Part of the peptide array showing peptides 33-48 corresponding to residues 160-250 of human 2N4R tau, incubated with recombinant BIN1-SH3-GST (top and bottom), or recombinant GST-only (middle), and probed with an antibody against BIN1 (middle and bottom) or no primary antibody (control, top). Black arrow shows positive signal at peptide 41 corresponding to amino acids 201-225 of human 2N4R tau. B) Human 2N4R tau sequence showing the proline-rich region of tau corresponding to peptide 41, which includes proline 216 (black arrow). C) HEK293 cells were transfected with wild type 2N4R tau (WT) or PxxP mutant tau constructs in which a single proline residue at site 176, 182, 200, 206, 213, 216, 219 or 223 was mutated to alanine to disrupt the PxxP sequence. Proteins in lysates from HEK293 cells (input) were pulled down with BIN1-SH3-GST beads, and western blotted with antibodies against total tau (top) GST (bottom). D) The amount of tau pulled down by GST was quantified and the bar chart shows this data relative to WT 2N4R tau (control). When P216 was mutated to alanine, tau binding to BIN1-SH3 was significantly reduced (*p<0.05). Following D’Agostino and Pearson normality testing, data were analysed using Kruskal-Wallis test and Dunn’s multiple comparisons test. Data shown are mean ± S.E.M, n=9.

Phosphorylation of tau at serine and threonine residues affects its interactions with other SH3-domain containing proteins (38). Tau in lysates from primary cortical neurons treated with 50 nM okadaic acid (OA), a protein phosphatase inhibitor (39, 40), for 4 hours to increase tau phosphorylation (Figure 3A), showed only trace amounts of binding to BIN1-SH3-GST (Figure 3B). In similar experiments using lysates from neurons treated with 25 mM of the glycogen synthase-3 inhibitor lithium chloride (LiCl) (39, 41) (Figure 3C), or 20 μM of the casein kinase-1 inhibitor IC261 (39, 42) to dephosphorylate tau (Supplementary Figure 3A), the amount of tau pulled down by BIN1-SH3 was similar to controls (Figure 3D, Supplementary Figure 3B). These data suggest that BIN1-tau interactions, mediated by BIN1-SH3 and ^213^PxxPxxP^219^ in tau, take place predominantly when tau is dephosphorylated.

**Figure 3:**
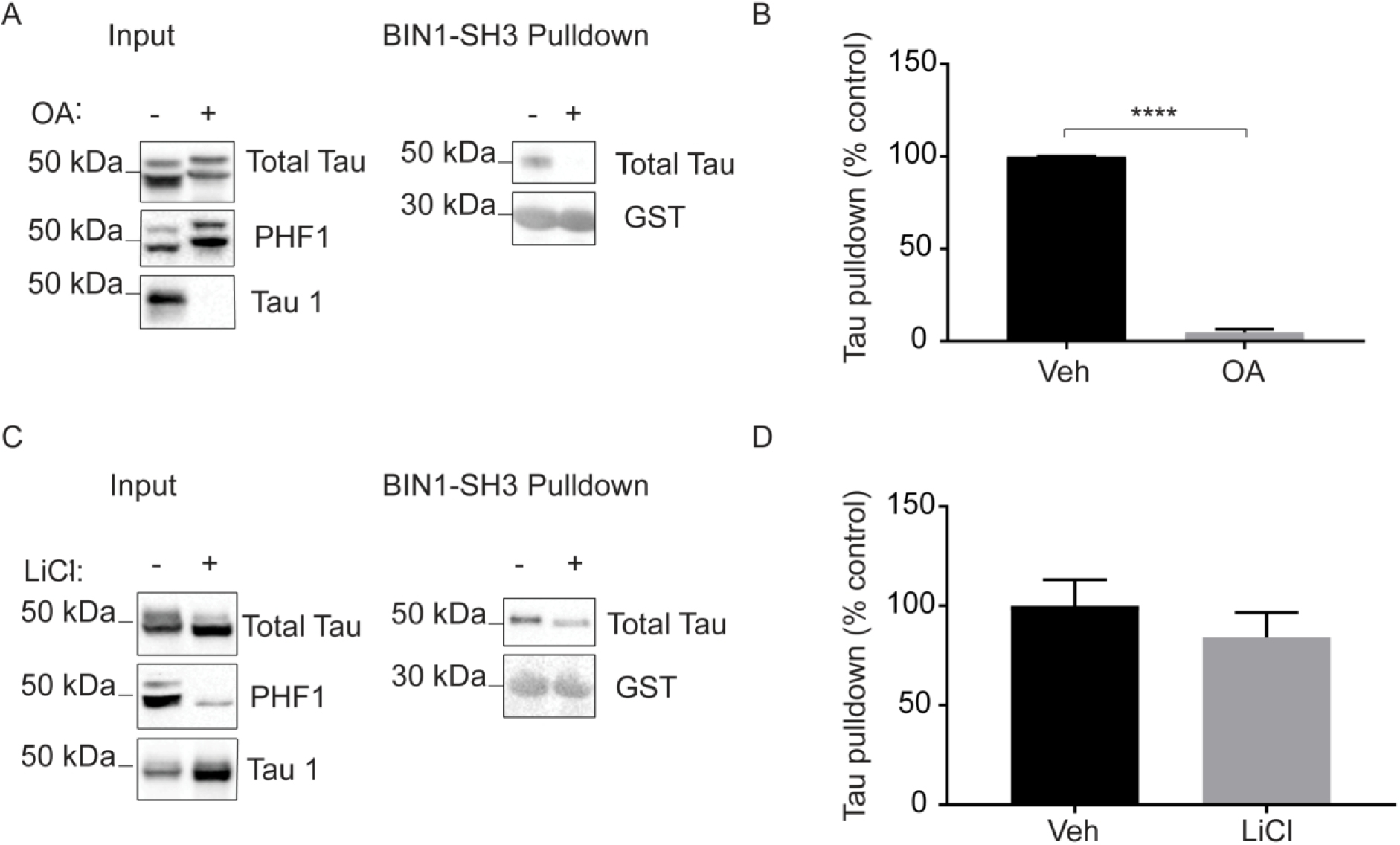
Increasing serine/threonine tau phosphorylation abolishes the interaction of tau with BIN1-SH3. A) Primary cortical neurons were treated with either vehicle (−) or 50 nM okadaic acid (OA,+) for 4 hours. Proteins were pulled down from lysates with BIN1-SH3-GST. Western blots of neuronal lysates (input) with antibodies against total tau (top), tau phosphorylated at Ser394/404 (PHF1, middle) and tau dephosphorylated at Ser199/202/Thr 205 (Tau-1, bottom) show increased tau phosphorylation following okadaic acid treatment. B) Quantification of the amount of tau from vehicle- or okadaic acid-treated neurons pulled down by BIN1-SH3-GST shown as percentage mean control (vehicle). The amount of tau pulled down by BIN1-SH3-GST was reduced following okadaic acid treatment of primary neurons. Following Shapiro-Wilk normality testing, the data were analysed using an unpaired T-test. Data shown are mean ± S.E.M, n=4. ****p<0.0001. C) Lysates from primary cortical neurons show reduced tau phosphorylation following treatment with 25 mM LiCl (+) for 4 hours relative to vehicle-treated neurons (-). BIN1-SH3-GST pulldowns show that there was no apparent difference in the amount of tau pulled down by BIN1-SH3-GST following LiCl treatment. D) Quantification of the amount of tau from vehicle- or LiCl-treated neurons pulled down by BIN1-SH3-GST shown as percentage mean control (vehicle). Following Shapiro-Wilk normality testing, data were analysed using a Mann-Whitney test. Data shown are mean ± S.E.M, n = 3.

### BIN1 loss in cytoplasmic fractions of AD brain correlates with increased synaptic tau

In AD brain, highly phosphorylated tau is mislocalised to synaptic compartments (31) where tau is believed to exert toxicity (43, 44), and there is a loss of the longest isoform of BIN1 (9, 10). We examined temporal cortex from control (Braak stage 0-III), moderate (Braak stage III-IV) and severe (Braak stage V-VI) post-mortem brain on western blots. In total brain homogenates, we confirmed a trend towards reduction of BIN1 in severe relative to moderate AD and control tissues (Figure 4A, B), and significant increases in total tau amounts with increasing disease severity (Figure 4A, C). Protein amounts were normalised to neuron-specific enolase in the same sample prior to quantification to account for any effects of neuronal loss and/or gliosis (45). Cytoplasmic and synaptoneurosome fractions (31) were isolated from the same brain samples (Supplementary Figure 4) and these showed a significant accumulation of tau in the synaptic compartment in severe AD, relative to moderate AD and controls (Figure 4D, F). Notably, the accumulation of synaptic tau paralleled the loss of cytoplasmic tau suggesting altered tau localisation (Figure 4H, J). There was also an accumulation of synaptic tau phosphorylated at S396/S404 (PHF1) in severe and moderate AD versus controls (Figure 4D, G). No differences in BIN1 in synaptoneurosomes were observed between groups (Figure 4D, E). However, there were significant decreases in cytoplasmic BIN1 with increasing disease stage (Figure 4H, I) that correlated positively with reductions in cytoplasmic tau (Figure 4J, 5A) and inversely with increased synaptic tau (Figure 5B, C). Taken together, these data suggest that loss of cytoplasmic BIN1 facilitates the mis-sorting of tau to the synapse.

**Figure 4:**
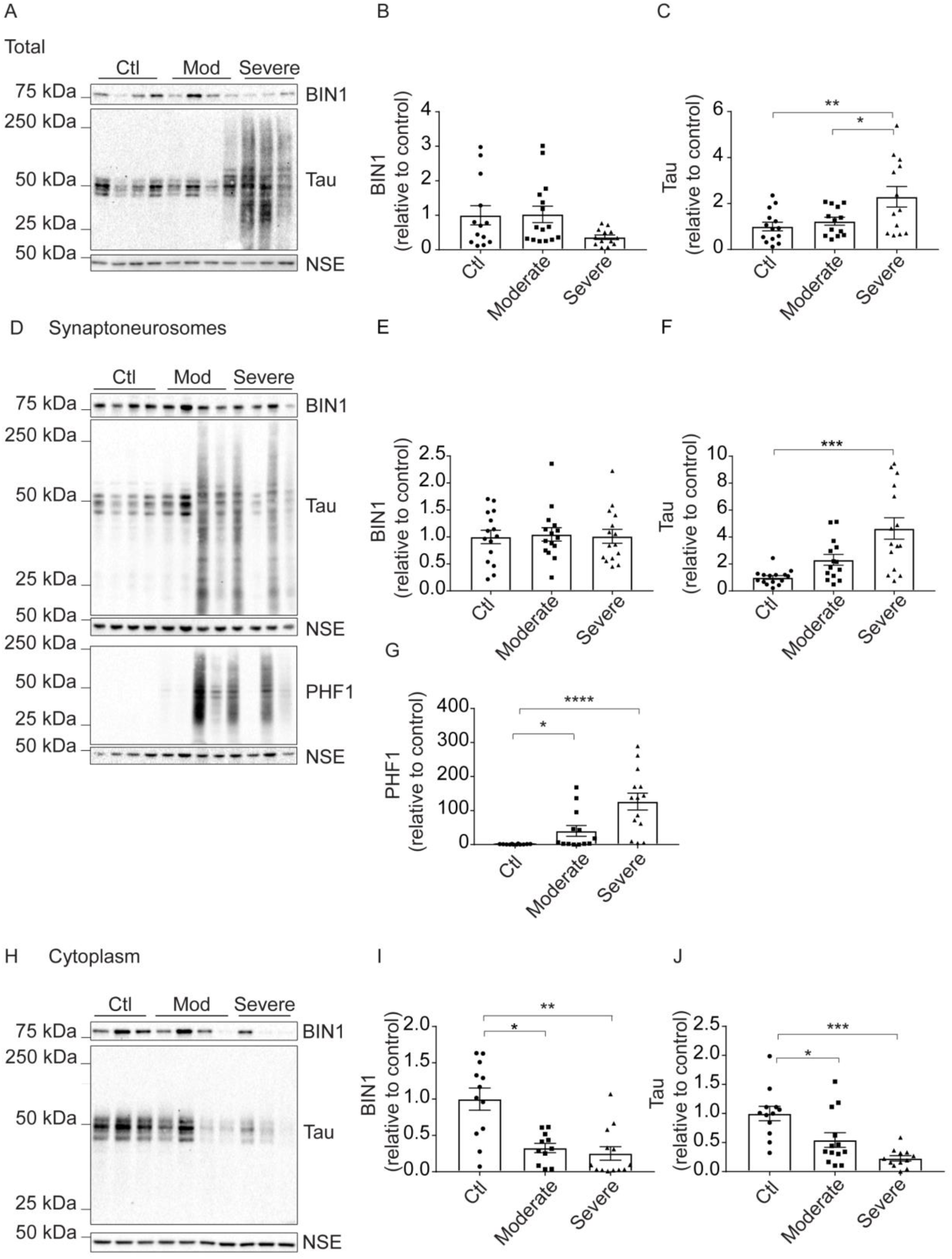
BIN1 and tau are reduced in the cytoplasm and phosphorylated tau accumulates in the synaptic fraction of AD temporal cortex. A) Total homogenates from temporal cortex homogenates of control (Braak stage 0-III), moderate (Braak stage III-IV) and severe (Braak stage V-VI) AD brain were western blotted using antibodies against BIN1 (top), total tau (middle), and neuron specific enolase (NSE, bottom). Bar charts show quantification of B) BIN1 and C) tau amounts following normalisation to NSE in the same sample. Data shown are mean ± S.E.M. expressed as a fold change of average control. Following D’Agostino and Pearson normality testing, data were analysed using a one-way ANOVA with Holm-Sidak’s multiple comparisons test. n=13 (BIN1) or 14 (tau). D) Synaptoneurosomes isolated from the same temporal cortex samples were also immunoblotted with antibodies against BIN1, tau, tau phosphorylated at Ser396/404 (PHF1), and neuron-specific enolase (NSE). Bar charts show quantification of E) BIN1, F) tau, and G) PHF1 in synaptoneurosomes following normalisation to NSE in the same sample. Data are mean ± S.E.M. expressed as a proportion of average control. Following D’Agostino and Pearson normality testing, data was analysed using non-parametric Kruskal-Wallis test with Dunn’s multiple comparison test. n=15 (BIN1 and tau) or 12 (PHF1). H) The cytoplasmic fraction was blotted as above with antibodies against BIN1, tau, and NSE. Bar charts show quantification of I) BIN1 and J) tau in the cytoplasmic fraction following normalisation to NSE in the same sample. Data are mean ± S.E.M. expressed as proportion mean control. Following D’Agostino and Pearson normality testing, BIN1 data was analysed using non-parametric Kruskal-Wallis test with Dunn’s multiple comparison test and tau data using a one-way ANOVA with Holm-Sidak’s multiple comparisons test. n=11 (BIN1) or 12 (tau). *p<0.05, ** p<0.01, ***p<0.001, ****p<0.0001.

**Figure 5:**
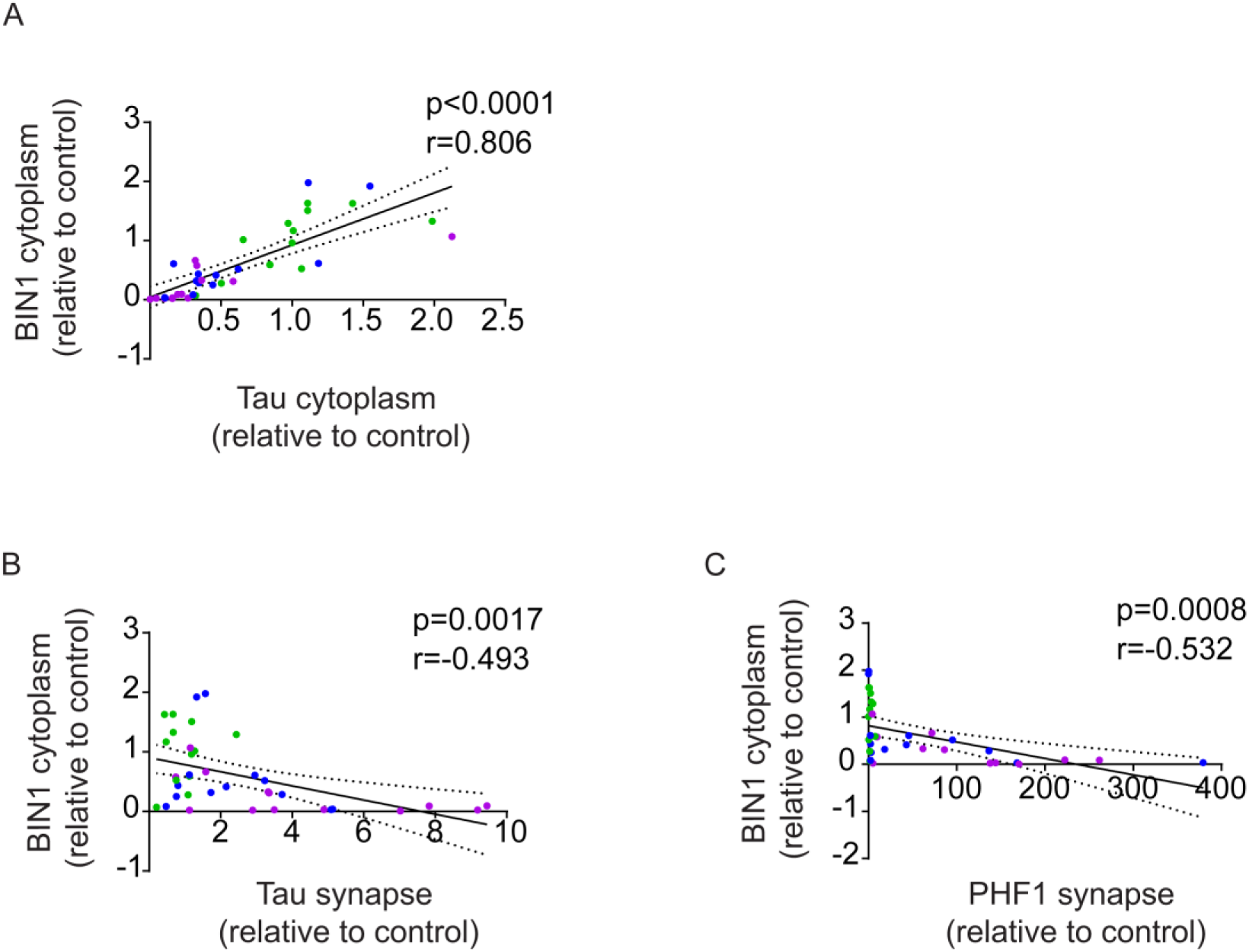
Loss of BIN1 correlates with loss of cytoplasmic tau and increased synaptic tau. Correlation analysis of BIN1 and tau amounts in A) cytoplasmic fractions shows a strong positive correlation between BIN1 and tau (n=38), and strong negative correlations between B) cytoplasmic BIN1 and synaptic tau (n=38), and C) cytoplasmic BIN1 and synaptic tau phosphorylated at Ser396/404 (PHF1) (n=36). Colours represent mild (green), moderate (blue) and severe (purple) stage samples.

### BIN1 knockdown causes synaptic accumulation of phosphorylated tau in neurons

To investigate the effects of lowering BIN1 expression on tau mislocalisation, we knocked down BIN1 in rat primary cortical neurons using lentivirus. In control neurons, BIN1 and tau are found in both cytoplasmic and synaptic fractions (Fig. 6A). N-SIM imaging showed that BIN1 decorates MAP2-positive and MAP2-negative fibres (Figure 6B) and localises in close proximity to pre-synaptic (synaptophysin) and post-synaptic (PSD95) markers (Figure 6C). BIN1 knockdown did not alter the total amount of tau or its phosphorylation status, or the amounts of PSD95 and synaptophysin in neuronal cell lysates (Supplementary Figure 5). However, BIN1 knockdown resulted in a significant increase in the amounts of tau phosphorylated at the PHF-1 epitope (phospho-Ser396/404) in synaptic fractions, relative to controls (Figure 6D, F). These data support the idea that reduced cytoplasmic BIN1 in AD brain could act in parallel with increased tau phosphorylation to enable mislocalisation of phosphorylated tau to synapses.

**Figure 6:**
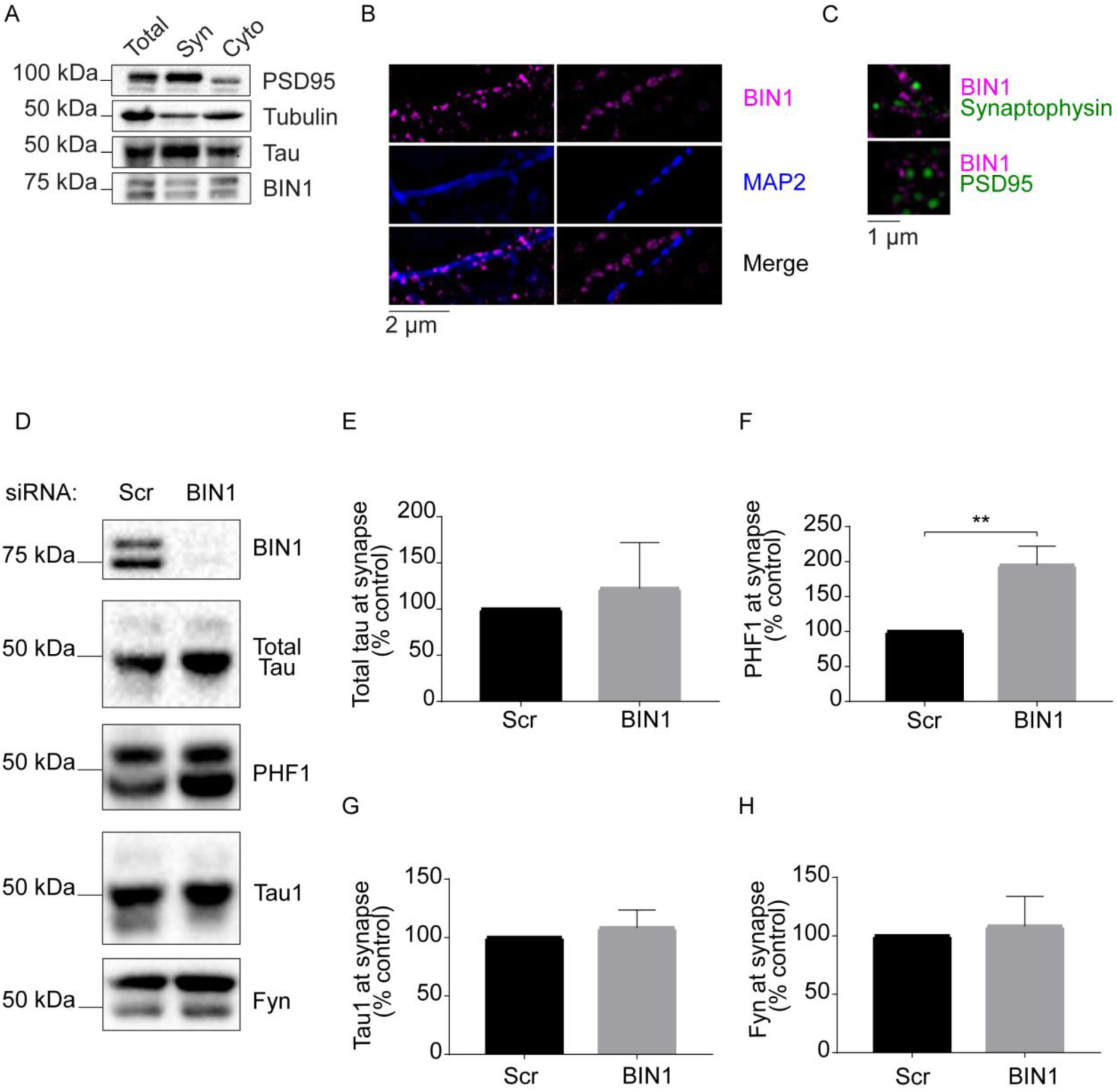
BIN1 is present at synapses and its knockdown increases the synaptic abundance of phosphorylated tau in primary neurons. A) 22 DIV primary cortical neurons biochemically fractionated into total, synaptic protein-enriched (syn) and cytoplasmic (cyto) fractions were western blotted with antibodies against PSD95, tubulin, tau and BIN1 and show the presence of BIN1 and tau in the synaptic fraction. B) N-SIM super resolution images of primary cortical neurons immunolabelled with antibodies against BIN1 (ab54764, pink) and the dendritic marker MAP2 (blue) showing that BIN1 is present within dendrites and axons in cultured neurons. C) N-SIM super-resolution images show a close association of BIN1 (ab54764, pink) with the pre-synaptic marker synaptophysin (top image, green) and the post-synaptic marker PSD95 (bottom image, green). D) Lysates from primary cortical neurons transduced with scrambled control shRNA (Scr) lentivirus or BIN1 shRNA (BIN1) lentivirus and biochemically fractionated as above were immunoblotted with antibodies against BIN1, total tau, tau phosphorylated at Ser396/404 (PHF1), tau dephosphorylated at Ser199/202/Thr205 (Tau-1) and Fyn. Western blots show quantification of synaptic protein amounts for E) tau, F) PHF1, G) Tau-1 and H) Fyn. Data were normalised to the synaptic marker PSD95 in the same sample and are expressed as percentage mean control (scrambled siRNA). Data are mean ± S.E.M. and were analysed using Mann-Whitney test. n=4-5, **p<0.01.

When bound to tau, the non-receptor tyrosine kinase, Fyn, is trafficked to the post-synapses where it is believed to mediate β-amyloid toxicity in AD (46). We previously showed that the Fyn-SH3 domain also binds preferentially to proline 216 in tau (26, 37). We therefore examined Fyn localisation in synaptic fractions following BIN1 knockdown. We found no alterations in synaptic Fyn in neurons treated with BIN1 siRNA (Figure 6H), suggesting that Fyn does not compete with BIN1 for binding to tau.

### Loss of BIN1 alters spine morphology and reduces tau release from neurons

Modulating BIN1 expression was recently shown to affect dendritic spine morphology and AMPA receptor-mediated synaptic transmission via changes in AMPA receptor surface expression and trafficking (22, 29). Since we and others have previously shown that neuronal depolarisation and stimulation of AMPA receptors mediates tau release (23, 27), we examined the effects of BIN1 knockdown on tau release. We first confirmed that BIN1 knockdown affects synaptic morphology in 23 DIV primary neurons exogenously expressing eGFP using iSIM. BIN1 knockdown did not cause any alterations in dendritic spine length, volume or density (Figure 7A-D), but resulted in significant increases in the diameter of spine heads and necks (Figure 7E-F) and a reduced head:neck diameter ratio (Figure 7G). When spine morphologies were examined, BIN1 knockdown had no effect on the proportion of immature stubby or thin spines (Figure 7H, I), but significantly reduced the proportion of filopodia (Figure 7K) and increased the proportion of mature mushroom spines (Figure 7J), which are relatively stable and have a high density of AMPA receptors (47–50).

**Figure 7:**
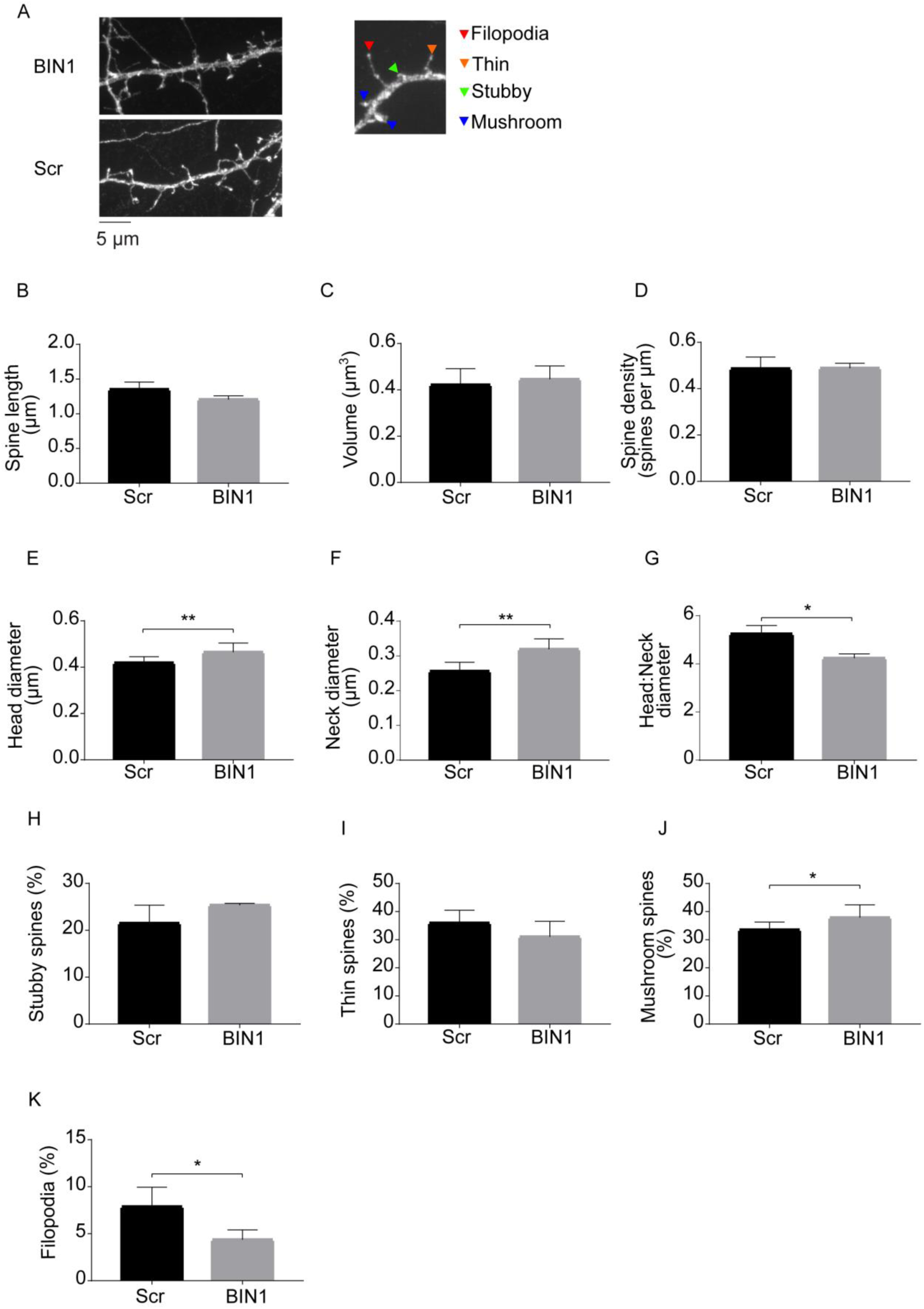
BIN1 knockdown alters dendritic spine morphology. A) Primary cortical neurons transduced with BIN1 shRNA (BIN1) lentivirus (top left) or scrambled control shRNA (Scr) lentivirus (bottom left) were transfected with a plasmid expressing eGFP and fixed at 23 DIV. Maximum intensity projections were generated from Z-stacks acquired using I-SIM super-resolution imaging. Dendritic spines were classified as either filopodia or stubby, thin or mushroom spines (right). Bar charts show quantification of spine B) length, C) volume, D) density, E) head diameter, F) neck diameter, G) ratio of spine head to neck diameter, and percentage of H) stubby, I) thin, J) mushroom spines and K) filopodia. Data are mean ± S.E.M. and were analysed using a randomised block 2-way ANOVA. n=3. *p<0.05, **p<0.01.

Tau release into culture medium from 21 DIV neurons was measured by ELISA and normalised to the total amount of tau in neurons from the same culture well (27). BIN1 knockdown caused a significant reduction in basal tau release without affecting the amount of intracellular tau (Figure 8A-C). BIN1 knockdown followed by depolarisation of neurons with KCl to stimulate tau release also significantly reduced tau release relative to controls (Figure 8D-F). The observed changes in tau release were not due to cell toxicity since there were no alterations in lactate dehydrogenase content in medium between conditions (Figure 8G). Thus, BIN1 knockdown results in alterations in synapse morphology and reduces basal and stimulated tau release.

**Figure 8:**
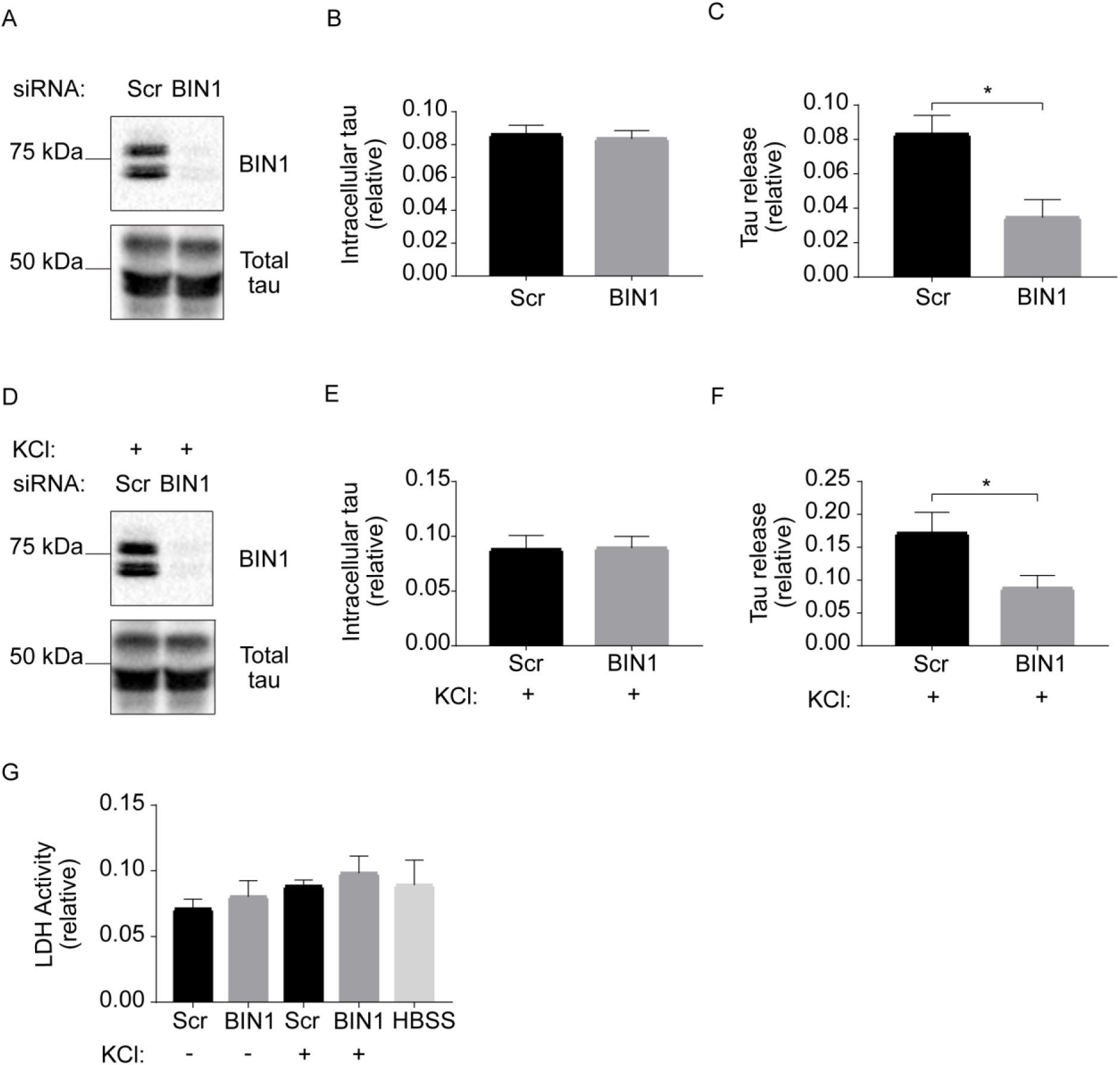
BIN1 knockdown reduces tau release. A) Cell lysates from 21 DIV primary cortical neurons transduced with scrambled control shRNA (Scr) lentivirus or BIN1 shRNA (BIN1) lentivirus were western blotted with antibodies against BIN1 and total tau. B) Quantification shows no effect of BIN1 knockdown on tau amount. Shapiro-Wilk normality test demonstrated that the data were normally distributed and data were analysed using an unpaired T-test. C) Tau content in conditioned media from neurons was determined by ELISA. Tau content in medium was quantified relative to intracellular tau levels and shows reduced tau release upon BIN1 knockdown. Shapiro-Wilk normality test demonstrated that the data were normally distributed, so data were analysed using an unpaired T-test. D) Cells transduced as above were depolarised with 50 nM KCl (+) for 30 minutes and the lysates were western blotted with antibodies against BIN1 and total tau. E) KCl treatment had no effect on intracellular tau amounts. Shapiro-Wilk normality test demonstrated that the data were normally distributed, so data were analysed using an unpaired T-test. F) Tau in conditioned media from KCl-stimulated cells was measured as described for basal conditions. Tau release from neurons in which BIN1 was knocked down remained reduced upon neuron depolarisation with KCl. Shapiro-Wilk normality test demonstrated that the data were not normally distributed, so data were analysed using a Mann-Whitney test. G) Lactate dehydrogenase amounts were measured in medium from unstimulated (-) or KCl-stimulated (+) primary cortical neurons transduced with scrambled control shRNA (Scr) lentivirus or BIN1 shRNA (BIN1) lentivirus and showed no effect of treatment on cell viability. Shapiro-Wilk normality test demonstrated that the data were not normally distributed, so data were analysed using a Kruskal-Wallis test with Dunn’s multiple comparison test. All graphs show mean ± S.E.M, n=7 (intracellular tau) or 6 (tau release, and lactate dehydrogenase assay). *p<0.05.

## Discussion

BIN1 is closely linked with tau abnormalities that underlie the progression of sporadic AD (12). Here, we have shown that BIN1 and tau interact via the SH3 domain of BIN1 and residues 201-225 in the proline rich region of tau. Notably, mutation of P216 to alanine, one of the seven PxxP motifs in tau, significantly reduces the interaction of these two proteins. Our data are in agreement with previous findings that residues 212-231 of tau were required for the interaction with BIN1-SH3 (20) further refining the site of interaction to include only residues 212-225 in tau. Malki et al. (18) also determined that a single site in tau likely interacts with a single BIN1 site on the basis of binding assays. Phosphorylation of tau at serine/threonine residues affects its interaction with other SH3 domain-containing proteins (26, 37, 38, 51), and we demonstrated that increasing tau phosphorylation in primary neurons strongly reduces tau-BIN1-SH3 binding. Our data, together with the fact that tau-phosphorylation by ERK prevents the interaction with the BIN1 SH3 domain (20), suggest that the increased phosphorylation of tau in AD would reduce the association of BIN1 with tau. This view is supported by the lack of BIN1 immunolabelling within tangle-bearing neurons in AD brain (13). However, others have shown co-localisation between BIN1 and neurofibrillary tangles in AD brain (11) and have reported the existence of a tau-BIN1-clusterin complex in AD brain (52). Our results suggest that changes in tau-BIN1 interactions may not be global but could occur primarily between BIN1 and largely dephosphorylated tau in neuronal cytoplasm. Increased tau phosphorylation in AD would disrupt tau binding to BIN1 and retention of tau in the cytoplasm, allowing phosphorylated tau to be mis-sorted to synapses.

Expression of the longest neuronal isoform of BIN1 is reported to be lost in AD (10). We have extended this characterisation here to show a significant loss of cytoplasmic BIN1 from Braak stage III onwards in AD brain tissue. From Braak stage III this loss of BIN1 also showed a strong positive correlation with tau and mislocalisation of total and phosphorylated tau to the synaptic compartment, which may reflect an unbound pool of tau being phosphorylated and mis-sorted. In support of this idea, increased expression of BIN1 increases tau-BIN1 interactions and reduces tau-tubulin interactions (19), which would affect tau localisation. This is an important concept since alterations in the trafficking and normal positioning of tau are considered to be early pathogenic changes in AD (53–57).

To mirror the loss of BIN1 in AD, we knocked down BIN1 in rat primary neurons and this resulted in an accumulation of phosphorylated tau at synapses. In parallel, over-expression of BIN1 in mice expressing human tau led to a reduced number of somatodendritic tau inclusions (19). While tau has been observed at both the pre- and post-synapse in control post-mortem brain, phosphorylated tau is generally only found at the synapse in AD (32) where it is associated with loss of synaptic function, dysregulated calcium signalling, aberrant trafficking of glutamate receptors, and loss of mushroom spines (46, 58, 59). It is tempting to speculate therefore, that loss of BIN1 could promote AD by allowing the accumulation of damaging forms of phosphorylated tau at the synapse.

We show here that BIN1 knockdown results in spines with a larger head and neck diameter and increases the ratio of more mature mushroom spines to immature filopodia. In agreement with this, BIN1 over-expression results in the opposite changes to spines, leading to structural alterations in the hippocampus and memory deficits (22, 60). Mushroom spines are considered to be more stable and in general as spine size increases the number of AMPA receptors on the spine increases, increasing synaptic strength (49, 61, 62). This may appear to be in contrast with our assertion that BIN1 knockdown promotes synapse damage by directing phosphorylated tau into synapses. However, synapse enlargement is reported to occur in the early stages of neurodegeneration in AD (63) as a compensatory mechanism for the pre-synapse loss occurring early in the disease process (64), which could explain the increased spine diameter we report here. It is also possible that the effect of altering BIN1 expression on dendritic spines is independent of its binding to tau. Schurmann *et al.* (29) report that BIN1 modulates trafficking from recycling endosomes to the cell surface thereby altering the surface localisation of AMPA receptors in dendritic spines (29). Hence alterations in vesicle trafficking may be another mechanism by which BIN1 alters the structure of dendritic spines.

Finally, we demonstrate that loss of BIN1 expression results in reduced tau secretion from neurons, both in basal conditions and following neuronal depolarisation with KCl. We previously characterised the tau released from primary neurons as being largely intact, dephosphorylated (23), and likely to reflect trans-cellular signalling functions of extracellular tau (65, 66), distinguishing it from the aggregated, cleaved and highly phosphorylated tau species implicated in trans-synaptic tau spread/propagation. The results in this study suggest that the function of intact dephosphorylated tau is disrupted when BIN1 is lost. However, several different mechanisms have been suggested to underlie tau secretion in health and disease (66, 67). Tau fragments containing the proline-rich region are enriched among secreted tau fragments (67) while those lacking the proline-rich region are not secreted (65). Our ELISA only allows measurement of tau containing the proline-rich region of tau, therefore we cannot rule out the possibility that the release of distinct tau fragments could also be affected by BIN1 knockdown. Nevertheless, we report a novel finding that lowering BIN1 levels, as occurs in AD, results in significant inhibition of tau release from wild-type rat neurons. This would doubtless affect trans-cellular signalling functions of tau, including that at muscarinic acetylcholine receptors (68).

Our data demonstrate that BIN1 binds tau in a phosphorylation-dependent manner and this interaction is mediated by the BIN1-SH3 and tau proline-rich domains. We find that BIN1 is reduced in AD cytoplasm, and that this correlates with the loss of cytoplasmic tau and its accumulation in synaptic fractions. Modelling the loss of BIN1 in AD in primary neurons showed that when BIN1 is knocked down, mislocalisation of phosphorylated tau to synaptic fractions is increased, in parallel with reduced tau secretion. We hypothesise therefore, that disruptions to BIN1-tau interactions caused by BIN1 loss in AD enables tau-mediated synaptotoxicity and disrupts extracellular tau functions. Our data provide a potential mechanism by which polymorphisms near *BIN1* may increase AD risk and/or progression.

## Acknowledgements

This work was supported by Alzheimer’s Research UK (ARUK-RF2015-5, ARUK-PhD2017-4 to EG and ARUK-RF2014-2 to BP-N and ARUK-EG2013-B1 to WN). We are grateful to Professor Isabelle Landrieu (University of Lille Nord de France) for her generous gift of BIN1-SH3 domain plasmid, and Professor Peter Davies (Feinstein Institute of Medical Research, NY, USA) for his kind gift of tau antibodies. Some of this data was previously published as abstracts and posters at the 14^th^ international conference on Alzheimer’s and Parkinson’s Diseases (AD/PD 2019) and at Alzheimer’s Research UK Conferences in 2017, 2018 and 2019.

## Disclosures

The authors declare no competing interests

## Author Contributions

EG performed most experiments and analysed data; MT, DL, CT, RG, CE, RK, DH and BP-N performed additional experiments, and provided expertise and advice. EG and WN designed the research and wrote the paper.

## REFERENCES

1. Hu X, Pickering E, Liu YC, Hall S, Fournier H, Katz E, et al. (2011): Meta-analysis for genome-wide association study identifies multiple variants at the BIN1 locus associated with late-onset Alzheimer’s disease. PLoS One. 6:e16616.

2. Naj AC, Jun G, Reitz C, Kunkle BW, Perry W, Park YS, et al. (2014): Effects of multiple genetic Loci on age at onset in late-onset Alzheimer disease: a genome-wide association study. JAMA neurology. 71:1394–1404.

3. Seshadri S, Fitzpatrick AL, Ikram MA, DeStefano AL, Gudnason V, Boada M, et al. (2010): Genome-wide analysis of genetic loci associated with Alzheimer disease. JAMA. 303:1832–1840.

4. Wijsman EM, Pankratz ND, Choi Y, Rothstein JH, Faber KM, Cheng R, et al. (2011): Genome-wide association of familial late-onset Alzheimer’s disease replicates BIN1 and CLU and nominates CUGBP2 in interaction with APOE. PLoS Genet. 7:e1001308.

5. Miyashita A, Koike A, Jun G, Wang LS, Takahashi S, Matsubara E, et al. (2013): SORL1 is genetically associated with late-onset Alzheimer’s disease in Japanese, Koreans and Caucasians. PLoS One. 8:e58618.

6. Reitz C, Jun G, Naj A, Rajbhandary R, Vardarajan BN, Wang LS, et al. (2013): Variants in the ATP-binding cassette transporter (ABCA7), apolipoprotein E 4,and the risk of late-onset Alzheimer disease in African Americans. JAMA. 309:1483–1492.

7. Lambert JC, Ibrahim-Verbaas CA, Harold D, Naj AC, Sims R, Bellenguez C, et al. (2013): Meta-analysis of 74,046 individuals identifies 11 new susceptibility loci for Alzheimer’s disease. Nat Genet. 45:1452–1458.

8. Prokic I, Cowling BS, Laporte J (2014): Amphiphysin 2 (BIN1) in physiology and diseases. J Mol Med (Berl). 92:453–463.

9. De Rossi P, Buggia-Prevot V, Clayton BL, Vasquez JB, van Sanford C, Andrew RJ, et al. (2016): Predominant expression of Alzheimer’s disease-associated BIN1 in mature oligodendrocytes and localization to white matter tracts. Mol Neurodegener. 11:59.

10. Glennon EB, Whitehouse IJ, Miners JS, Kehoe PG, Love S, Kellett KA, et al. (2013): BIN1 is decreased in sporadic but not familial Alzheimer’s disease or in aging. PLoS One. 8:e78806.

11. Holler CJ, Davis PR, Beckett TL, Platt TL, Webb RL, Head E, et al. (2014): Bridging integrator 1 (BIN1) protein expression increases in the Alzheimer’s disease brain and correlates with neurofibrillary tangle pathology. J Alzheimers Dis. 42:1221–1227.

12. Chapuis J, Hansmannel F, Gistelinck M, Mounier A, Van Cauwenberghe C, Kolen KV, et al. (2013): Increased expression of BIN1 mediates Alzheimer genetic risk by modulating tau pathology. Mol Psychiatry. 18:1225–1234.

13. Adams SL, Tilton K, Kozubek JA, Seshadri S, Delalle I (2016): Subcellular Changes in Bridging Integrator 1 Protein Expression in the Cerebral Cortex During the Progression of Alzheimer Disease Pathology. J Neuropathol Exp Neurol.

14. Bungenberg J, Surano N, Grote A, Surges R, Pernhorst K, Hofmann A, et al. (2016): Gene expression variance in hippocampal tissue of temporal lobe epilepsy patients corresponds to differential memory performance. Neurobiology of disease. 86:121–130.

15. De Rossi P, Buggia-Prevot V, Andrew RJ, Krause SV, Woo E, Nelson PT, et al. (2017): BIN1 localization is distinct from Tau tangles in Alzheimer’s disease. Matters (Zur). 2017.

16. Bejanin A, Schonhaut DR, La Joie R, Kramer JH, Baker SL, Sosa N, et al. (2017): Tau pathology and neurodegeneration contribute to cognitive impairment in Alzheimer’s disease. Brain : a journal of neurology. 140:3286–3300.

17. Guo T, Dakkak D, Rodriguez-Martin T, Noble W, Hanger DP (2019): A pathogenic tau fragment compromises microtubules, disrupts insulin signaling and induces the unfolded protein response. Acta Neuropathol Commun. 7:2.

18. Malki I, Cantrelle FX, Sottejeau Y, Lippens G, Lambert JC, Landrieu I (2017): Regulation of the interaction between the neuronal BIN1 isoform 1 and Tau proteins - role of the SH3 domain. The FEBS journal. 284:3218–3229.

19. Sartori M, Mendes T, Desai S, Lasorsa A, Herledan A, Malmanche N, et al. (2019): BIN1 recovers tauopathy-induced long-term memory deficits in mice and interacts with Tau through Thr(348) phosphorylation. Acta neuropathologica.

20. Sottejeau Y, Bretteville A, Cantrelle FX, Malmanche N, Demiaute F, Mendes T, et al. (2015): Tau phosphorylation regulates the interaction between BIN1’s SH3 domain and Tau’s proline-rich domain. Acta Neuropathol Commun. 3:58.

21. Wang HF, Wan Y, Hao XK, Cao L, Zhu XC, Jiang T, et al. (2016): Bridging Integrator 1 (BIN1) Genotypes Mediate Alzheimer’s Disease Risk by Altering Neuronal Degeneration. J Alzheimers Dis. 52:179–190.

22. Daudin R, Marechal D, Wang Q, Abe Y, Bourg N, Sartori M, et al. (2018): BIN1 genetic risk factor for Alzheimer is sufficient to induce early structural tract alterations in entorhinal cortex-dentate gyrus pathway and related hippocampal multi-scale impairments. bioRxiv.437228.

23. Pooler AM, Phillips EC, Lau DH, Noble W, Hanger DP (2013): Physiological release of endogenous tau is stimulated by neuronal activity. EMBO Rep. 14:389–394.

24. Gomez-Suaga P, Perez-Nievas BG, Glennon EB, Lau DHW, Paillusson S, Morotz GM, et al. (2019): The VAPB-PTPIP51 endoplasmic reticulum-mitochondria tethering proteins are present in neuronal synapses and regulate synaptic activity. Acta Neuropathol Commun. 7:35.

25. Hill RE, Eaton-Rye JJ (2014): Plasmid construction by SLIC or sequence and ligation-independent cloning. Methods Mol Biol. 1116:25–36.

26. Lau DH, Hogseth M, Phillips EC, O’Neill MJ, Pooler AM, Noble W, et al. (2016): Critical residues involved in tau binding to fyn: implications for tau phosphorylation in Alzheimer’s disease. Acta Neuropathol Commun. 4:49.

27. Croft CL, Wade MA, Kurbatskaya K, Mastrandreas P, Hughes MM, Phillips EC, et al. (2017): Membrane association and release of wild-type and pathological tau from organotypic brain slice cultures. Cell death & disease. 8:e2671.

28. Schindelin J, Arganda-Carreras I, Frise E, Kaynig V, Longair M, Pietzsch T, et al. (2012): Fiji: an open-source platform for biological-image analysis. Nat Methods. 9:676–682.

29. Schurmann B, Bermingham DP, Kopeikina KJ, Myczek K, Yoon S, Horan KE, et al. (2019): A novel role for the late-onset Alzheimer’s disease (LOAD)-associated protein Bin1 in regulating postsynaptic trafficking and glutamatergic signaling. Mol Psychiatry.

30. Frandemiche ML, De Seranno S, Rush T, Borel E, Elie A, Arnal I, et al. (2014): Activity-dependent tau protein translocation to excitatory synapse is disrupted by exposure to amyloid-beta oligomers. J Neurosci. 34:6084–6097.

31. Perez-Nievas BG, Stein TD, Tai HC, Dols-Icardo O, Scotton TC, Barroeta-Espar I, et al. (2013): Dissecting phenotypic traits linked to human resilience to Alzheimer’s pathology. Brain : a journal of neurology. 136:2510–2526.

32. Tai HC, Serrano-Pozo A, Hashimoto T, Frosch MP, Spires-Jones TL, Hyman BT (2012): The synaptic accumulation of hyperphosphorylated tau oligomers in Alzheimer disease is associated with dysfunction of the ubiquitin-proteasome system. Am J Pathol. 181:1426–1435.

33. Byrne AM, Elliott C, Hoffmann R, Baillie GS (2015): The activity of cAMP-phosphodiesterase 4D7 (PDE4D7) is regulated by protein kinase A-dependent phosphorylation within its unique N-terminus. Febs Letters. 589:750–755.

34. Correas I, Padilla R, Avila J (1990): The tubulin-binding sequence of brain microtubule-associated proteins, tau and MAP-2, is also involved in actin binding. Biochem J. 269:61–64.

35. Drager NM, Nachman E, Winterhoff M, Bruhmann S, Shah P, Katsinelos T, et al. (2017): Bin1 directly remodels actin dynamics through its BAR domain. EMBO Rep. 18:2051–2066.

36. He HJ, Wang XS, Pan R, Wang DL, Liu MN, He RQ (2009): The proline-rich domain of tau plays a role in interactions with actin. BMC Cell Biol. 10:81.

37. Usardi A, Pooler AM, Seereeram A, Reynolds CH, Derkinderen P, Anderton B, et al. (2011): Tyrosine phosphorylation of tau regulates its interactions with Fyn SH2 domains, but not SH3 domains, altering the cellular localization of tau. The FEBS journal. 278:2927–2937.

38. Reynolds CH, Garwood CJ, Wray S, Price C, Kellie S, Perera T, et al. (2008): Phosphorylation regulates tau interactions with Src homology 3 domains of phosphatidylinositol 3-kinase, phospholipase Cgamma1, Grb2, and Src family kinases. J Biol Chem. 283:18177–18186.

39. Pooler AM, Usardi A, Evans CJ, Philpott KL, Noble W, Hanger DP (2012): Dynamic association of tau with neuronal membranes is regulated by phosphorylation. Neurobiol Aging. 33:431 e427–438.

40. Van Dolah FM, Ramsdell JS (1992): Okadaic acid inhibits a protein phosphatase activity involved in formation of the mitotic spindle of GH4 rat pituitary cells. J Cell Physiol. 152:190–198.

41. Stambolic V, Ruel L, Woodgett JR (1996): Lithium inhibits glycogen synthase kinase-3 activity and mimics wingless signalling in intact cells. Current biology : CB. 6:1664–1668.

42. Behrend L, Milne DM, Stoter M, Deppert W, Campbell LE, Meek DW, et al. (2000): IC261, a specific inhibitor of the protein kinases casein kinase 1-delta and -epsilon, triggers the mitotic checkpoint and induces p53-dependent postmitotic effects. Oncogene. 19:5303–5313.

43. McInnes J, Wierda K, Snellinx A, Bounti L, Wang YC, Stancu IC, et al. (2018): Synaptogyrin-3 Mediates Presynaptic Dysfunction Induced by Tau. Neuron. 97:823–835 e828.

44. Zhou L, McInnes J, Wierda K, Holt M, Herrmann AG, Jackson RJ, et al. (2017): Tau association with synaptic vesicles causes presynaptic dysfunction. Nat Commun. 8:15295.

45. Kurbatskaya K, Phillips EC, Croft CL, Dentoni G, Hughes MM, Wade MA, et al. (2016): Upregulation of calpain activity precedes tau phosphorylation and loss of synaptic proteins in Alzheimer’s disease brain. Acta Neuropathol Commun. 4:34.

46. Ittner LM, Ke YD, Delerue F, Bi M, Gladbach A, van Eersel J, et al. (2010): Dendritic function of tau mediates amyloid-beta toxicity in Alzheimer’s disease mouse models. Cell. 142:387–397.

47. Bourne J, Harris KM (2007): Do thin spines learn to be mushroom spines that remember? Current opinion in neurobiology. 17:381–386.

48. Dorostkar MM, Zou C, Blazquez-Llorca L, Herms J (2015): Analyzing dendritic spine pathology in Alzheimer’s disease: problems and opportunities. Acta neuropathologica. 130:1–19.

49. Lee KF, Soares C, Beique JC (2012): Examining form and function of dendritic spines. Neural Plast. 2012:704103.

50. Segal M (2010): Dendritic spines, synaptic plasticity and neuronal survival: activity shapes dendritic spines to enhance neuronal viability. Eur J Neurosci. 31:2178–2184.

51. Lee G, Newman ST, Gard DL, Band H, Panchamoorthy G (1998): Tau interacts with src-family non-receptor tyrosine kinases. J Cell Sci. 111 (Pt 21):3167–3177.

52. Zhou Y, Hayashi I, Wong J, Tugusheva K, Renger JJ, Zerbinatti C (2014): Intracellular clusterin interacts with brain isoforms of the bridging integrator 1 and with the microtubule-associated protein Tau in Alzheimer’s disease. PLoS One. 9:e103187.

53. Braak H, Braak E (1991): Neuropathological stageing of Alzheimer-related changes. Acta neuropathologica. 82:239–259.

54. Braak H, Thal DR, Ghebremedhin E, Del Tredici K (2011): Stages of the pathologic process in Alzheimer disease: age categories from 1 to 100 years. J Neuropathol Exp Neurol. 70:960–969.

55. Frost B, Jacks RL, Diamond MI (2009): Propagation of tau misfolding from the outside to the inside of a cell. J Biol Chem. 284:12845–12852.

56. Zempel H, Mandelkow E (2014): Lost after translation: missorting of Tau protein and consequences for Alzheimer disease. Trends in neurosciences.

57. Zempel H, Thies E, Mandelkow E, Mandelkow EM (2010): Abeta oligomers cause localized Ca(2+) elevation, missorting of endogenous Tau into dendrites, Tau phosphorylation, and destruction of microtubules and spines. J Neurosci. 30:11938–11950.

58. Hoover BR, Reed MN, Su J, Penrod RD, Kotilinek LA, Grant MK, et al. (2010): Tau mislocalization to dendritic spines mediates synaptic dysfunction independently of neurodegeneration. Neuron. 68:1067–1081.

59. Tracy TE, Gan L (2018): Tau-mediated synaptic and neuronal dysfunction in neurodegenerative disease. Current opinion in neurobiology. 51:134–138.

60. Sartori M, Mendes T, Desai S, Lasorsa A, Herldean A, Malmanche N, et al. (2018): BIN1 recovers tauopathy-induced long-term memory deficits in mice and interacts with Tau through Thr348 phosphorylation. bioRxiv.

61. Hanley JG (2008): AMPA receptor trafficking pathways and links to dendritic spine morphogenesis. Cell Adh Migr. 2:276–282.

62. Woolfrey KM, Srivastava DP (2016): Control of Dendritic Spine Morphological and Functional Plasticity by Small GTPases. Neural Plast. 2016:3025948.

63. DeKosky ST, Scheff SW (1990): Synapse loss in frontal cortex biopsies in Alzheimer’s disease: correlation with cognitive severity. Annals of neurology. 27:457–464.

64. Serrano-Pozo A, Frosch MP, Masliah E, Hyman BT (2011): Neuropathological alterations in Alzheimer disease. Cold Spring Harb Perspect Med. 1:a006189.

65. Perez M, Cuadros R, Hernandez F, Avila J (2016): Secretion of full-length tau or tau fragments in a cell culture model. Neurosci Lett. 634:63–69.

66. Perez M, Medina M, Hernandez F, Avila J (2018): Secretion of full-length Tau or Tau fragments in cell culture models. Propagation of Tau in vivo and in vitro. Biomol Concepts. 9:1–11.

67. Kanmert D, Cantlon A, Muratore CR, Jin M, O’Malley TT, Lee G, et al. (2015): C-Terminally Truncated Forms of Tau, But Not Full-Length Tau or Its C-Terminal Fragments, Are Released from Neurons Independently of Cell Death. J Neurosci. 35:10851–10865.

68. Gomez-Ramos A, Diaz-Hernandez M, Rubio A, Miras-Portugal MT, Avila J (2008): Extracellular tau promotes intracellular calcium increase through M1 and M3 muscarinic receptors in neuronal cells. Mol Cell Neurosci. 37:673–681.

