## Supplemental data for "Loss of the Alzheimer’s-linked bridging integrator 1 (BIN1) protein affects synaptic structure and disrupts tau localisation and release"

**Supplementary Figure 1**

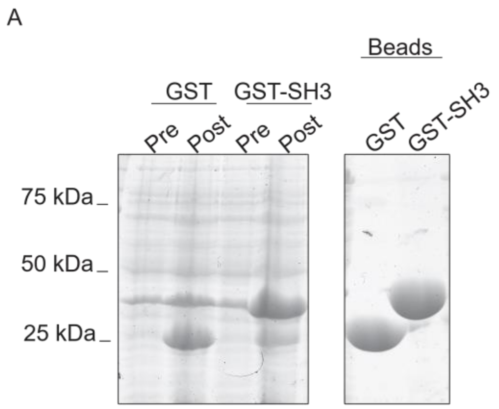


**Supplementary Figure 1: Preparation of GST-BIN1-SH3 beads.** A) Coomassie blue-stained SDS-PAGE gel showing GST-only and BIN1-SH3-GST-expressing *E.coli* cultures pre- and post- induction of protein expression (left) and BIN1-SH3-GST and GST purified on glutathione-S-sepharose beads.

**Supplementary Figure 2**

A

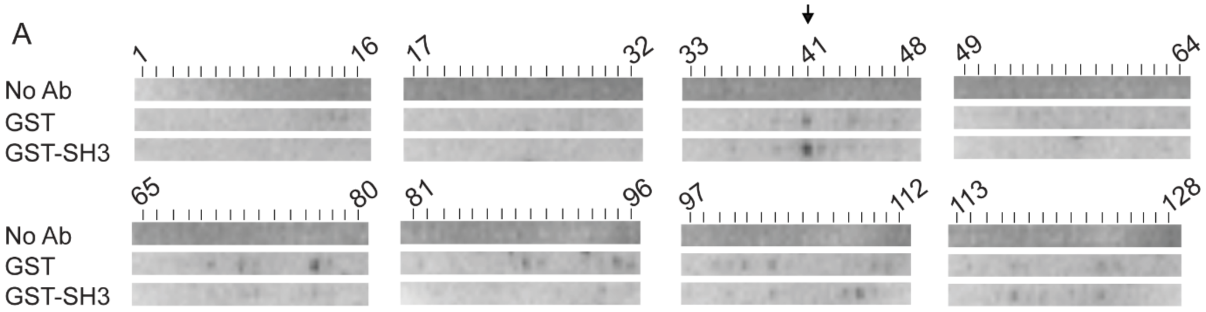


B

151 IATPRGAAPP GQKGQANATR IPAKT**p**PAPK T**p**PSSGEPPK SGDRSGYSS**p** 200
201 GSPGT**p**GSRS RT**p**SL**p**TP**p**T REPKKVAVVR TP**p**KSPSSAK SRLQTAPVPM 250

**Supplementary Figure 2: The BIN1-SH3 domain binds to residues 201-225 in tau.** Peptide array consisting of consecutive and overlapping 25 amino acid peptides sequences of human 2N4R tau (1-85), the alternative splice site for 2N3R tau (86-94) and as a negative control the astrocytic protein S100B (95-124), all incubated with recombinant BIN1-SH3-GST (top and bottom), or recombinant GST only (middle), then probed with anti-BIN1 antibody 99D (middle and bottom) or no primary antibody control (top). Black arrow indicates peptide 41. B) Amino acid sequence 151-250 of human 2N4R tau in which the seven PxxP motifs are underlined, and prolines in these motifs that were mutated to alanine are indicated in lowercase and bold type.

**Supplementary Figure 3**

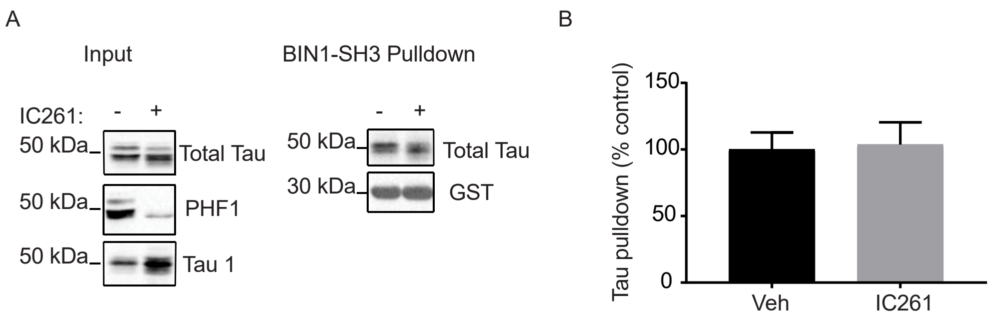


**Supplementary Figure 3: Inhibiting casein-kinase 1 activity to reduce tau phosphorylation does not alter the interaction of tau with BIN1-SH3.** A) Lysates from 21-23 DIV primary cortical neurons show reduced tau phosphorylation following treatment with 20 µM IC261 (+) for 4 hours relative to vehicle-treated neurons (-). BIN1-SH3-GST pulldowns show that there was no apparent difference in the amount of tau pulled down by BIN1-SH3-GST following reduction of tau phosphorylation by IC261 treatment. B) Quantification of the amount of tau from vehicle- or IC261-treated neurons pulled down by BIN1-SH3-GST shown as percentage mean control (vehicle). Following Shapiro-Wilk normality testing, data were analysed using a Mann-Whitney test. Data is mean ± S.E.M., n = 3.

**Supplementary Figure 4**

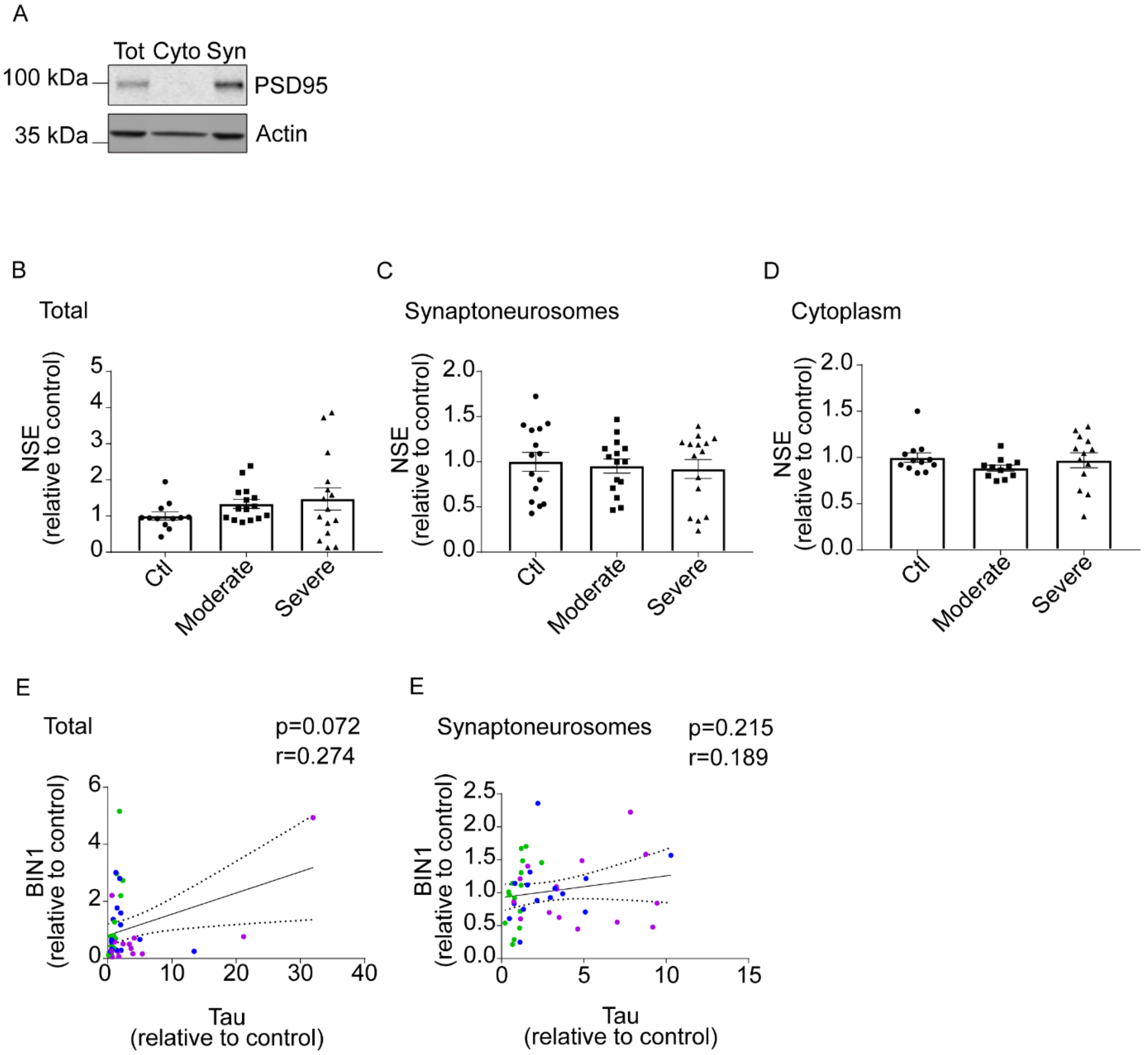


**Supplementary Figure 4: Fractionation of AD temporal cortex** A) Total temporal cortex homogenates and cytoplasmic and synaptoneurosome fractions isolated from the same brain samples were western blotted using antibodies against PSD95 (top), and actin (bottom) loading to confirm that synaptic proteins are found within the synaptoneurosome but not cytosolic fraction. Total homogenates, synaptoneurosome and cytoplasmic fractions of control (Braak stage 0-III), moderate (Braak stage III-IV) and severe (Braak stage V-VI) AD brain were immunoblotted and probed for neuron-specific enolase (NSE) (Figure 4). Bar charts show quantification of NSE in B) total (n=13), C) synaptoneurosome (n=15) and D) cytoplasmic fractions (n=11) following normalisation to controls.

Following D'Agostino and Pearson normality testing, data were analysed by Kruskal-Wallis test with Dunn's multiple comparison test (total and cytoplasm) or one-way ANOVA with Tukey's multiple comparisons test (synaptoneurosomes). Graphs show mean ± S.E.M. Correlation analysis of BIN1 and tau amounts in E) total homogenates (n=44), and F) synaptoneurosomes (n=45) shows no correlation between tau and BIN1 in these fractions. Colours in E and F represent mild (green), moderate (blue) and severe (purple) stage samples.

**Supplementary Figure 5**

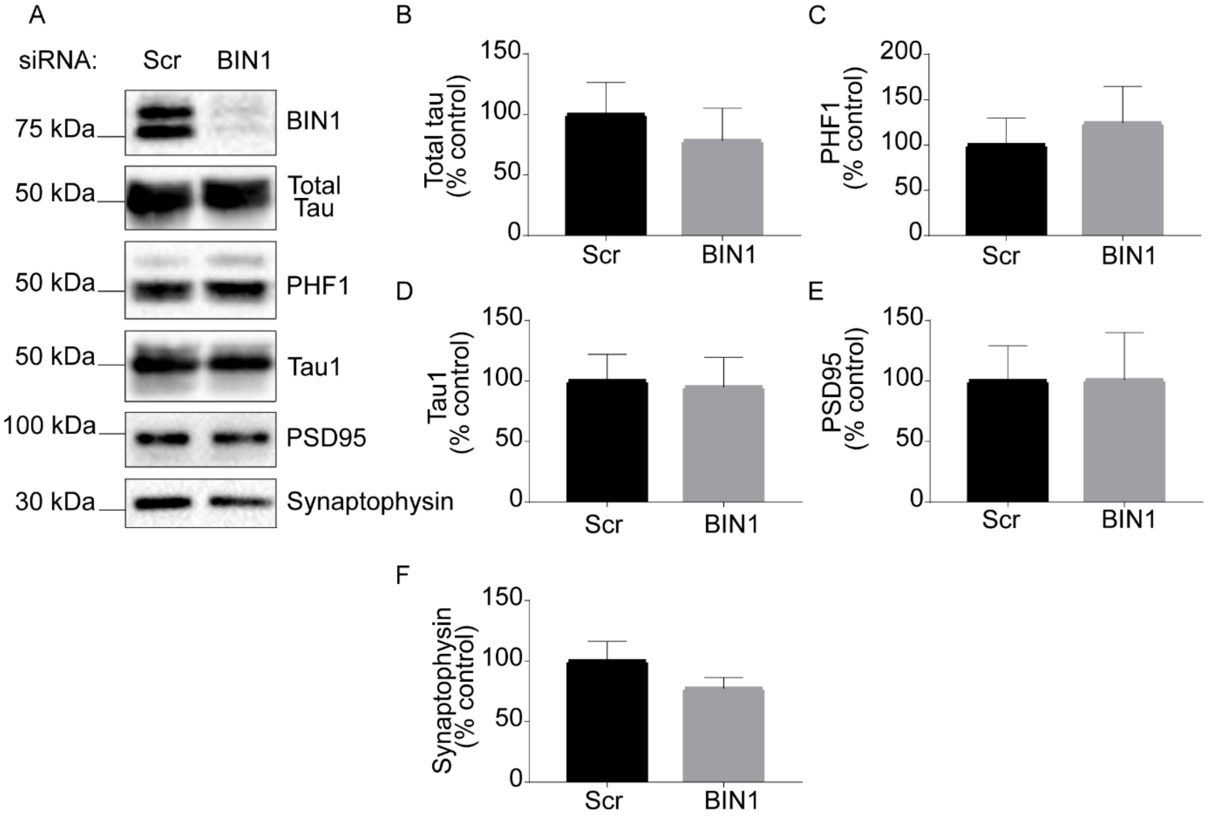


**Supplementary Figure 5: BIN1 knockdown does not alter tau or synaptic markers in cortical neurons.** A) Lysates from primary cortical neurons transduced with scrambled control shRNA (Scr) lentivirus or BIN1 shRNA (BIN1) lentivirus were western blotted for BIN1, total tau, tau phosphorylated at Ser396/404 (PHF1), tau dephosphorylated at Ser199/202/Thr205 (tau-1), PSD95, and synaptophysin. Quantification of western blots for B) total tau, C) PHF1, D) tau 1, E) PSD95, and F) synaptophysin. Data are expressed as a percentage of average control (Scr). Following Shapiro-Wilk normality testing total tau, tau1, PSD95, and synaptophysin were analysed using an un-paired T-test. PHF1 data were analysed by Mann-Whitney test. Graphs show the mean ± S.E.M of 3 (total tau, PHF1 tau and tau 1), 7 (PSD95) or 6 (synaptophysin) independent experiments.
